# Image-based force inference by biomechanical simulation

**DOI:** 10.1101/2023.12.01.569682

**Authors:** Michiel Vanslambrouck, Wim Thiels, Jef Vangheel, Casper van Bavel, Bart Smeets, Rob Jelier

## Abstract

During morphogenesis, cells precisely generate forces that drive cell shape changes and cellular motion. These forces predominantly arise from contractility of the actomyosin cortex, allowing for cortical tension, protrusion formation, and cell division. Image-based force inference can derive such forces from microscopy images, without complicated and time-consuming experimental set-ups. However, current methods do not account for common effects, such as physical confinement and local force generation. Here we propose a force-inference method based on a biophysical model of cell shape, and assess relative cellular surface tension, adhesive tension between cells, as well as cytokinesis and protrusion formation. We applied our method on fluorescent microscopy images of the early *C. elegans* embryo. Predictions for cell surface tension at the 7-cell stage were validated by measurements using cortical laser ablation. Our non-invasive method facilitates the accurate tracking of force generation, and offers many new perspectives for studying morphogenesis.

**Author summary:** An important challenge in understanding morphogenesis is to determine where forces are generated. Force generation, such as differential cortical tension, plays a key role in cellular self-organization. However, measuring these forces experimentally is a big challenge that typically requires complicated experiments and direct access to the cells being investigated. A non-invasive method, such as force inference based on cell shapes, would have important advantages. Unfortunately, current methods can only be applied in certain cases. Here, we describe a more flexible 3D force inference method, FIDES, that can account for physical confinement, cell divisions and protrusions. We applied the method to infer forces throughout early *C. elegans* embryogenesis. By using cortical laser ablations, we confirmed that FIDES correctly inferred cell surface tensions in a dynamic stage of the nematode’s embryogenesis. Our approach offers a route to routinely infer force generation in complex movements during morphogenesis, with microscopy images of cell shapes as the sole experimental input.

## Introduction

Understanding how cellular self-organization gives rise to functional multicellular systems is a central challenge in developmental biology. An important part of studying this process is characterizing the dynamic generation of force by the cells, as this highlights which active contributions need to be explained. Many of the involved forces are generated by the actomyosin cortex, which is a thin, highly dynamic mesh of actin fibers and myosin motors just underlying the cell membrane (Lecuit et al., 2011). The cell cortex can control cell shape by the generation of contractile forces. Contractility in the cortex, for instance, generates cortical tension, which causes cells to round up. The cortex can also give rise to specific structures, such as lamellipodia involved in cell migration, as well as the cytokinetic ring during cell division. Furthermore, the cortex attaches to adhesion receptors like cadherins, which are crucial for force transmission, cell-cell contacts, and cell shape. Cadherins can induce a reduction of actomyosin contractility in the cortex, enlarging the cell-cell contact area (Winklbauer, 2015; Maître et al., 2012). During morphogenesis, all these components work together to shape cells, perform divisions and ensure correct cellular positioning.

Many methods have been developed to measure the mechanical properties and force generation of biological systems, but most of them involve complicated and restrictive experimental procedures. For instance, methods such as atomic force microscopy and micropipette aspiration require a direct contact with the cell, which often requires the cell to be exposed outside its biophysical microenvironment (Lee and Liu, 2014; Haase and Pelling, 2015). Likewise, 3D traction force microscopy enables the measurement of how cells exert forces on their surroundings but involves embedding the cells in a gel (Legant et al., 2010). Finally, advanced experimental methods such as optical or magnetic tweezers (Zhang and Liu, 2008; Vlaminck and Dekker, 2012), FRET tension sensors (Freikamp et al., 2017), and laser ablations (Mayer et al., 2010), while powerful, demand challenging experimentation and tend to have limited throughput.

A promising alternative is to use cell shapes to infer the forces that shape them (Keller, 2013; Roffay et al., 2021), as microscopic cell imaging is relatively straightforward and non-invasive. One family of methods, referred to as Foam Force Inference (FFI) in this paper, assumes that the mechanics controlling cell shape are similar to that of droplets or bubbles in a foam. The shape of a foam is driven by surface tensions, which in the analogy match the tensions generated by the actomyosin contractility in the cell cortex. In FFI, the relative membrane tensions are inferred by measuring the contact angles between membranes at triple junctions. By assuming mechanical equilibrium at these triple junctions, the contact angles between the cells are determined by the relative surface tensions (Veldhuis et al., 2017; Xu et al., 2018) (eq. 2). Solving the set of equations corresponding to the observed triple junctions yields a surface tension for every cell-medium interface, and an interfacial tension for every specific cell-cell contact. Further, some FFI approaches incorporate the curvature of the membrane, by assuming that it is a function of the pressure differential across the membrane and the surface tension, following the Young–Laplace equation (see Materials and methods, eq. 1).

Several closely related algorithms have been developed since the original articles of Ishihara and Sugimura (2012) and Chiou et al. (2012), but all share the same assumptions about cell properties: Every cell has one uniform pressure and cell-medium surface tension, every cell-cell contact has a uniform interfacial tension, and cells behave like a foam in quasi-static mechanical equilibrium. Early implementations only considered 2D microscopy slices (Ishihara and Sugimura, 2012; Chiou et al., 2012; Brodland et al., 2014), but more recently, research shifted to 3D analysis (Veldhuis et al., 2017; Xu et al., 2018; Ichbiah et al., 2023). There are different ways to solve this problem, depending on the tissue structure and the cell properties that are inferred (Roffay et al., 2021). When both junction angles and curvatures are evaluated, we will call this curved FFI, which can infer both tensions and pressures. Meanwhile, tangent FFI considers only the junction angles, and does not allow relative pressures to be estimated. There has been some experimental validation for FFI. Veldhuis et al. (2017) inferred cell surface tensions in 8-cell mouse embryos and validated the results using micropipette aspiration. In *Drosophila*, Kong et al. (2019) used FFI to infer junction tensions in epithelium tissue and ommatidia, and validated the results with laser ablation of single junctions and groups of cells. More recently, Ichbiah et al. (2023) used FFI to infer forces in mouse and ascidian embryos, demonstrating that the inferred interfacial tensions explain observed cell shapes without the need for direct measurements. They also applied their method on *C. elegans* and discussed how the eggshell induces confinement, which is not accounted for by current implementations of FFI, though quasi-static foam models can account for such effects (Brakke, 1992). Indeed, despite the validation for passive non-constrained systems, the accuracy of FFI methods is likely to be affected by phenomena that are common in developing tissues, such as physical confinement, cell divisions, and protrusions. A notable example of a developing tissue that is not in equilibrium, is the 7-cell *C. elegans* embryo, on which Xu et al. (2018) applied curved FFI. During this specific developmental stage, assumptions of a fully passive foam model may be violated, as the embryo features active behaviors, including a dividing cell, several protrusions, and cell movements (Pohl and Bao, 2010).

Some of the limitations associated with FFI can be addressed by phase-field models, which use differential equations to model phase transitions rather than sharp interfaces. Kuang et al. (2022) performed phase-field simulations on early *C. elegans* embryos to estimate cell adhesion properties. The model can approximate various mechanical aspects, such as eggshell characteristics, volume conservation, and tension within a system. It is a novel technique for modeling cell behavior, and requires validation and establishing the bounds of its successful application.

Here, we introduce a simulation-based force inference system, called Force Inference by Discrete Element method Simulation (FIDES), which offers a more expressive model than classical FFI. Our approach initiates simulations directly from high-quality cell shape segmentations (Thiels et al., 2021), closely connecting force inferences with experimental microscopy data. Cells are represented using triangulated meshes, and forces are derived from the Deformable Cell Model (DCM), a mechanical model of cell shape solved numerically via the Discrete Element Method (Odenthal et al., 2013). This framework allows representing variations to cell surface tensions and cell-to-cell adhesion properties in a way similar to the models underlying FFI, while using a more flexible setup. Moreover, we incorporate an eggshell that confines the cells, as well as representations of local force generation to capture common active cellular behaviors. This includes a model for cell division to reproduce the cytokinetic furrow, which builds on a previous effort to capture cell division in the DCM framework (Cuvelier et al., 2023). Further, a simple protrusion model is used to represent lamellipodia-like shapes generated by the cells to probe their environment and exert forces on neighboring cells. As such, FIDES describes the embryo as a foam with active force models (Kim et al., 2021; Hallatschek et al., 2023), in contrast to FFI, which assumes a passive foam, without processes like protrusions. Optimization of model parameters is achieved such that simulated cells match observed cell shapes. This concept has some similarity to Noll et al. (2020), where model parameters were optimized to match image signal directly. In this article, we will compare FIDES to FFI implementations in the context of early *C. elegans* embryos, a well-known model for studying morphogenesis due to its transparency, small size, and nearly invariant development (Sulston et al., 1983). Direct measurement of cytoskeletal forces in *C. elegans* cells is challenging due to the eggshell that envelops the embryo. As part of the validation process, we will compare the inferred cell surface tensions with measurements obtained through cortical laser ablations. Additionally, we will evaluate our inferences over time and compare them with earlier research.

## Results

### Framework of FIDES

In our pipeline, we first employ confocal fluorescent microscopy to image cell membranes in *C. elegans* embryos. Each cell is subsequently segmented using the SDT-PiCS procedure (Thiels et al., 2021), and a single cell surface mesh is generated, as illustrated in fig. 1A. During the last step of this segmentation procedure, we perform a mechanical simulation where the mesh is attracted to the membrane signal, improving segmentation quality. The constraints posed by the DCM guarantee smooth cell shapes and realistic contact boundaries. The idea behind the FIDES method is to perform new simulations and extend them with realistic physical forces to stabilize the cell shapes without the membrane pixel attraction that was present during segmentation. This approach assumes that the observed cell shapes are close to mechanical equilibrium, which seems reasonable in early *C. elegans* development, where the influence of viscous forces is limited (Cuvelier et al., 2023).

**Fig 1.**
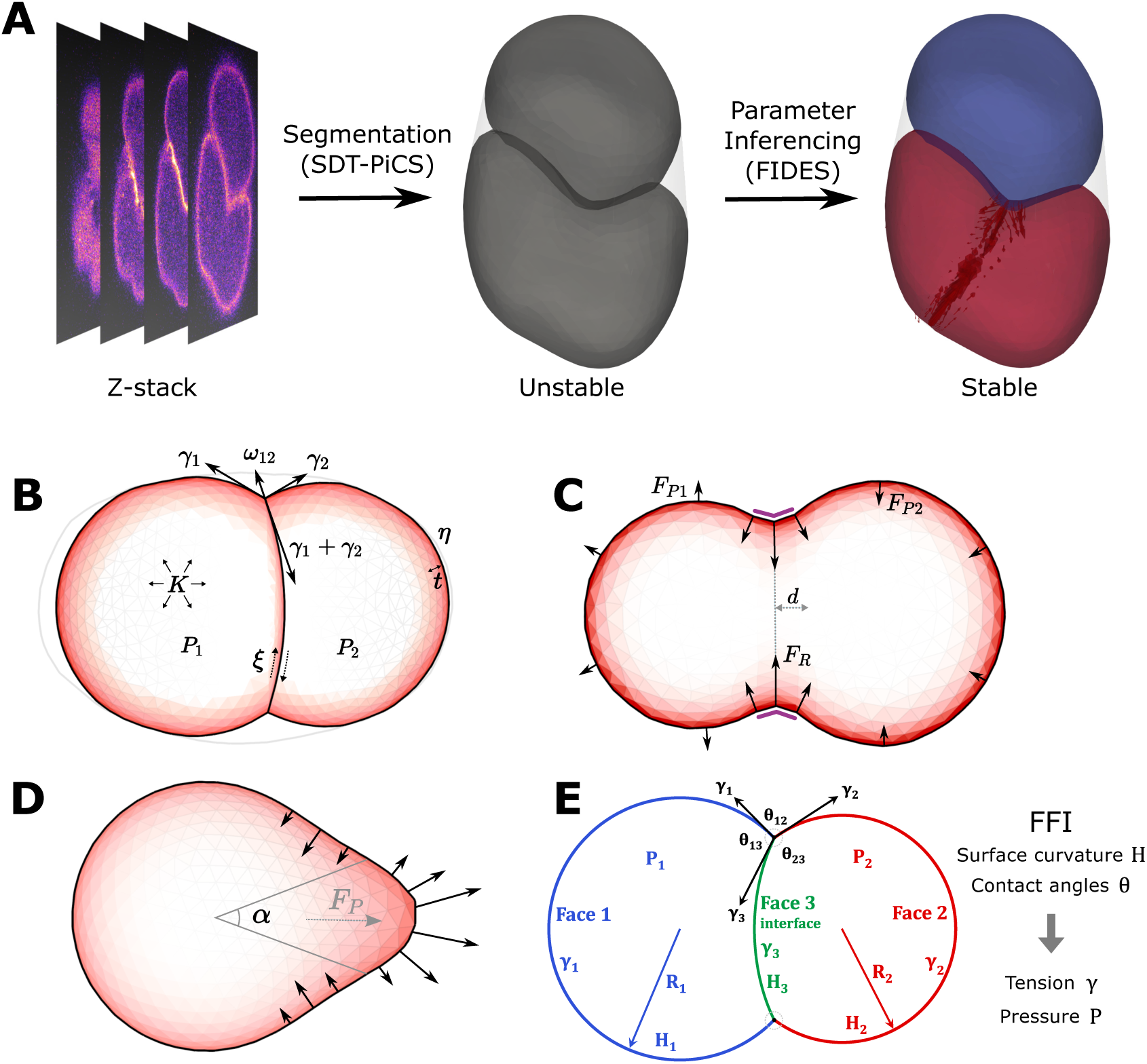
Overview of the FIDES pipeline and FFI. A: Pipeline for force inference. Confocal microscopy time-lapses are segmented using SDT-PiCS into 3D meshes of cells. The cell shapes serve as a starting point for DCM simulations and FIDES finds the optimal parameters that stabilize the cell shapes. In this example, a difference in surface tension is established, while the bottom cell also has a contractile ring. B: Description of the DCM. Single cells are represented as a triangulated shell, with surface tension *γ*, pressure *P*, viscosity *η*, cortical thickness *t*, and volume conservation regulated by bulk modulus *K*. Between cells, adhesive tension *ω* and friction *ξ* applies. C: Force model for cytokinetic ring. A ring of elastic springs is applied on a cell, which results in inward forces and creates the furrow. Two spherical coordinates determine the normal vector for the division plane, while distance *d* controls its offset from the mass center. An elastic spring constant *k* results in a ring force *F_R_*, which is distributed over neighboring nodes, making the furrow more smooth. The position of the ring is stabilized using an area modulus that resists changes in cell area on both sides of the ring with pressure force *F_P i_*. D: Force model for protrusion. The direction of the protrusion is controlled by two spherical coordinates, pointing from the mass center of the cell. The angle *α* controls the width of the protrusion and *F_P_* controls the total pushing force. The forces are smoothly distributed over the mesh according to a Difference of Gaussian kernels, with an outward force in the center of the protrusion, while the edges are pulled inwards to ensure an overall net force of zero. E: Overview of curved FFI. By assuming that cells behave like soap bubbles, we can write a series of equations that give observable variables, the contact angles *θ* at triple membrane junctions and the curvature *H* of the membranes, as a function of tensions *γ* and pressures *P* . This system of equations is next parametrized with measurements from segmented images and solved by least squares fitting.

We performed the embryo simulations with the Mpacts simulation framework, using the DCM model (fig. 1B) to approximate how cells take their shape (Odenthal et al., 2013). The setup permits modeling cell aggregates with full control over force generation and transmission. Specifically, it permits the assignment of different surface tensions to individual cells and a unique adhesive tension per cell-cell contact. In this paper, we use surface tension to refer to the cell-medium tension of a cell, whereas adhesive and interfacial tension specifically describe the interface between two cells. We consider surface tension *γ_i_* to be the tension on the free surface of a cell, while interfacial tension *γ_ij_* consists of two surface tensions, reduced by the adhesive tension *ω_ij_*. The adhesive tension can be seen as the reduction of cortical tension at the interface, for example following the effect of adhesion proteins on cortical contractility at the cell-cell interface (Arslan et al., 2021). Supplementary Text 1.1 describes the implementation of tension, pressure, as well as friction and viscous forces in the DCM. FIDES requires a complete parametrization of the model, where most of the physical parameters are fixed during the procedure (Supplementary Table S3).

To account for the variable cell shapes that are observed during embryogenesis, we added two models for local force generation. First, we modeled cell division, given that this is a frequent event during development. While mitotic rounding can be modeled as an increased cortical tension (Taubenberger et al., 2020), the cytokinetic furrow is produced by local contractile forces. Figure 1C shows how we model the furrow by selecting a ring of nodes along the cell mesh, and apply spring forces to constrict the furrow. To resist volume instability in the dividing cell, we introduce a bulk modulus that applies pressure to both daughters, similar to Cuvelier et al. (2023). Second, a protrusion model was introduced to accommodate cell shapes with lamellipodia. Cells are deformed by protrusive forces that originate from the localized polymerization of actin filaments (Lecuit et al., 2011). To simulate the formation of protrusions, we generate an outward force smoothly distributed over a portion of the cell’s surface, accompanied by an equal opposing force applied to the sides of the protrusion, using a Difference of Gaussians function to distribute the forces (fig. 1D). This way, our protrusion model acts as a localized force dipole. Both the cytokinetic ring and the protrusion model are minimally parametrized, defining mainly orientation and force (see Materials and methods). Finally, the eggshell is modeled as a stiff convex hull delimiting the embryo.

During the optimization procedure we fit a DCM model on the observed cell shapes. Cells are initialized with uniform parameters for surface tension *γ*, adhesive tension *ω*, and corresponding equilibrium volume *V^∗^*. The local force models are added where needed, each with several parameters that will be optimized. Next, we run the simulation and measure the error using the sum of displacements for the mesh nodes (see Materials and methods). Parameter inferencing uses stochastic optimization (Spall, 2011), where for every parameter, we randomly adjust its value and run a new simulation, accepting the change only if it improves the error. This procedure is repeated until the optimization converges, or the maximum number of iterations is reached.

FIDES yields the optimal surface tension, adhesive tension, and pressure parameters for the embryo, as well as descriptions of the cytokinetic rings and protrusions. The inferred tensions are relative values, as is the case for FFI.

Next, we assess the consistency of FIDES and the sensitivity to the chosen physical parameters. To this end, we performed a sensitivity analysis on three different embryos (Supplementary Figure S3). The results indicate that repeated experiments are highly consistent, with minimal variability. Further, the inferences are only minimally affected by changes in friction, viscosity, initial surface-to-adhesive tension ratio, or the simulation time step. However, a larger effect was observed for mesh size; when the triangles became too large, the loss of mesh detail led to less well-defined cell contacts that impacted the results.

### Comparison of FIDES to FFI on artificial data

In this section, we assess the performance of the FIDES pipeline and compare it to two implementations of FFI using several synthetic embryos. These synthetic embryos were generated using the DCM and augmented with our local force models, while varying the degree of complexity and realism. The objective is to validate the FIDES pipeline in ideal circumstances, but also to assess the sensitivity of FFI predictions to violations of the underlying assumptions. Specifically, foam models assume that cells are in quasi-static mechanical equilibrium, and that cell shapes are solely determined by isotropic cell-medium surface tension and interfacial tension.

To perform curved FFI, we reproduced the method of Xu et al. (2018) to work on our data. For every interface we calculate the average mean curvature *H*, and for every triple junction we sample the average contact angle *θ*. The Young-Laplace and Young-Dupré equations are then used to estimate the pressures, surface tensions and interfacial tensions between cells (fig. 1E). Our implementation for tangent FFI doesn’t use the curvatures and the Young-Laplace equations, and solves the system for only the tensions.

We start with a 4-cell synthetic embryo in a diamond configuration (fig. 2A), which adheres to the assumptions of foam models, and has its surface and interfacial tension parameters set arbitrarily. The model is similar to the real 4-cell *C. elegans* embryo, but lacks eggshell confinement. In this scenario, all methods demonstrate excellent predictive performance when comparing inferred and set values for surface and interfacial tension, with tangent FFI and FIDES achieving a Pearson’s r of 1.00 (fig. 2B).

**Fig 2.**
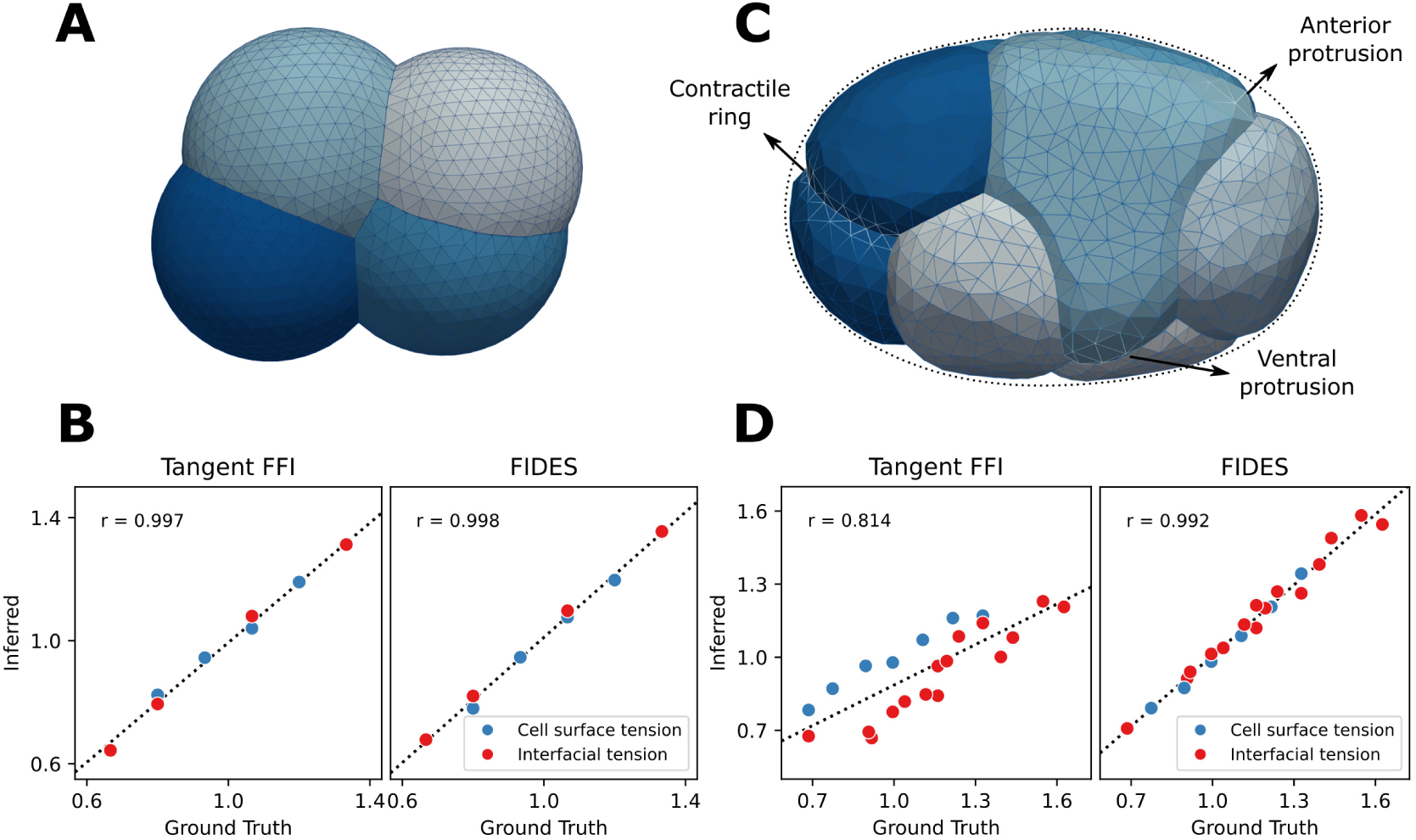
Performance of FFI and FIDES on synthetic embryos. A: Simple synthetic embryo, consisting of four cells having different volumes and surface tensions, and five unique cell-cell contacts with varying interfacial tension. B: Performance comparison of FFI and FIDES on the simple synthetic embryo. Both have an excellent correlation with the ground truth. C: Advanced synthetic embryo, modeled after the 7-cell *C. elegans* embryo. This sample features seven differently parametrized cells, 15 contacts with varying interfacial tension, a contractile ring on P2 and a total of four protrusions on ABpl and ABpr, two of which are visible. D: Performance comparison on the advanced synthetic embryo.

Next, we designed a more challenging situation, shown in fig. 2C, by attempting a reconstruction of the 7-cell *C. elegans* embryo, closely encapsulated in an eggshell, with a contractile ring on the P2 cell, a protrusion on ABpr and three protrusions on ABpl. These modifications make the synthetic embryo less quasi-static foam-like and could cause errors in FFI. First, encapsulation in an eggshell affects membrane curvatures and flattens triple junction angles at the embryo exterior. Second, local forces can change the geometry of contacts, both on the interior and exterior. For example, we observed that during cytokinesis, the contact between the dividing cell and other cells often takes on the shape of a saddle point, adding conflicting curvature information for curved FFI. In the case of the 7-cell synthetic embryo, tangent FFI now performs worse with a Pearson’s r of 0.81, while FIDES still performs well, achieving a score of 0.99 (fig. 2D). The results for curved FFI are worse than tangent FFI for the 7-cell case, with r = 0.68, as shown in Supplementary Figure S4. To see why FFI performed worse on this example, we next evaluated the FFI performance effects of local force models and eggshell confinement separately. Removing local forces from the example embryo only had a minor effect on the score of either curved or tangent FFI (table 1). For a synthetic 7-cell embryo without an eggshell, eliminating confinement, tangent FFI improved its score a lot. With both the eggshell removed and the local force models disabled, both tangent and curved FFI achieve accurate results.

**Table 1.** Contribution of factors disrupting FFI methods. The first case is the example displayed in fig. 2C, while the others are generated the same way, but with one or more features disabled. Pearson’s r values for correlation between method inferences and ground truth are made in the same way as in fig. 2.

| Condition | Tangent FFI | Curved FFI |
| --- | --- | --- |
| Confinement, local forces | 0.81 | 0.68 |
| Confinement, no local forces | 0.79 | 0.73 |
| No confinement, local forces | 0.95 | 0.76 |
| No confinement, no local forces | 0.98 | 0.96 |

For the remainder of the article, we will mainly focus on the better performing tangent FFI (see Discussion). Conversely, we can investigate how important these features of FIDES are in explaining the synthetic shapes. Keeping all features on the 7-cell synthetic embryo, but removing local force models or the eggshell from FIDES, leads to a drop in performance to 0.78 and 0.70, respectively.

### Validation of cortical tension inferences

To validate the results from force inference in vivo, we measured cell surface tension in the *C. elegans* embryo using cortical laser ablation experiments. Briefly, the cortical actomyosin mesh of a cell was imaged using confocal fluorescence microscopy in a *C. elegans* strain with fluorescently labelled actin filaments (F-actin) and non-muscular myosin (NMY-2). The cortex was then cut using a pulsed UV-laser, and the recoil movement of the cortex was quantified from the images, as illustrated in fig. 3A. Following Mayer et al. (2010), we treat the cortex as a viscoelastic material under contractile stress. When the cortex is cut, the tension is released and the cortex recoils following an exponentially decaying velocity profile (see Materials and methods, eq. 4). Assuming constant dampening, the exponential decay rate is then proportional to the cortical stiffness. Meanwhile, the initial recoil velocity is proportional to the cortical tension, and can then be used to validate image-based inferred surface tension forces. We recovered the recoil speeds from the images by manually tracking the movement of cortical highlights (fig. 3B). After combining all measurements per condition, an exponential decay function was fitted to the recoil speeds using non-linear least squares. The uncertainty on the exponential fit parameters was subsequently assessed using a bootstrap strategy (see Materials and methods).

**Fig 3.**
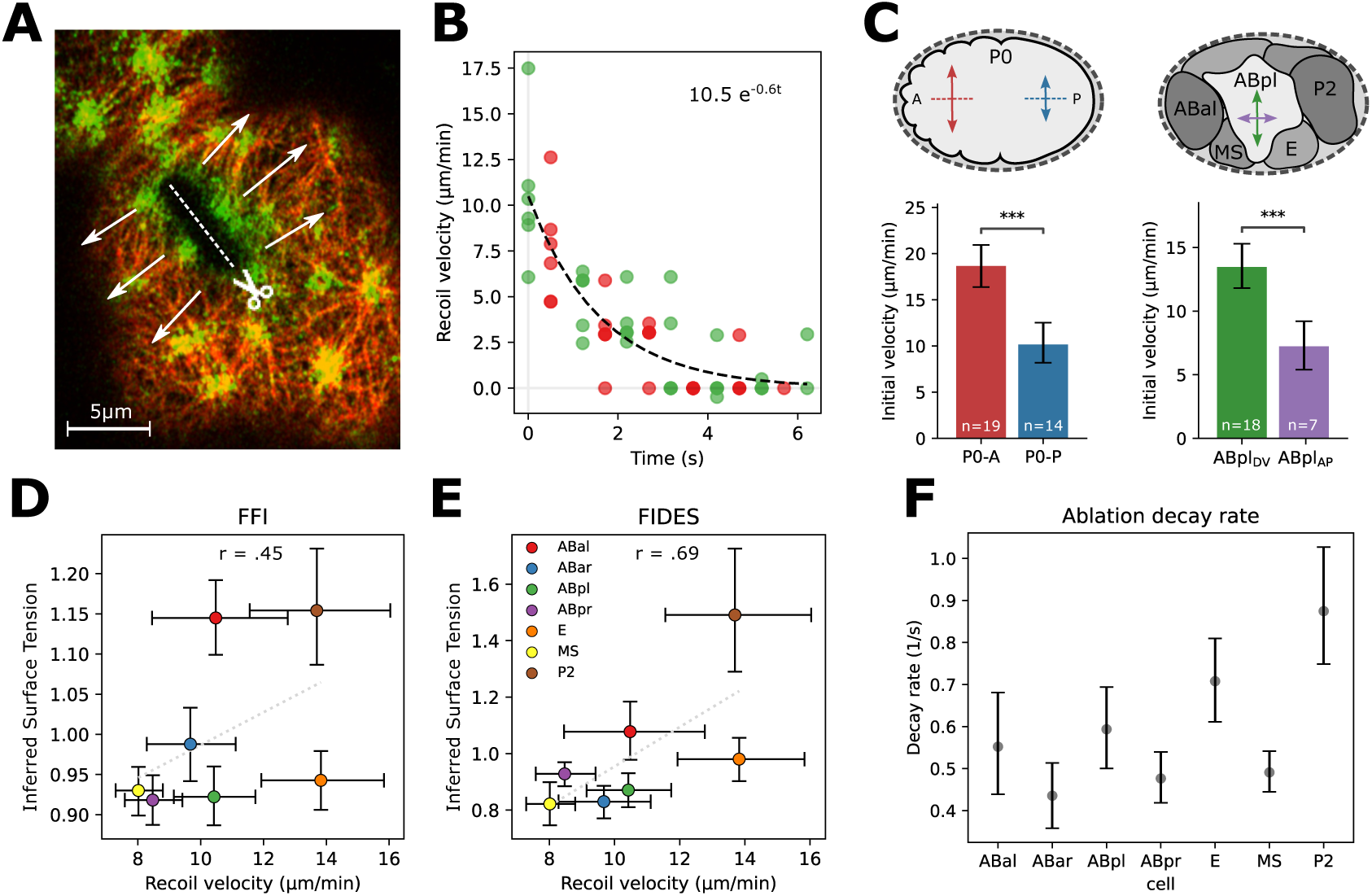
Cortical laser ablation as validation for cell surface tension inferences. A: Cortical laser ablation procedure for cell P0. A cut measuring 8 µm is made in the cortex, causing the cortex to recoil perpendicularly to the cut. Here, this cut happens in the anterior side of the polarizing zygote (P0-A). B: Highlights of the cortex are tracked over the duration of the ablation experiment, resulting in velocity measurements for the myosin marker (green) and the actin marker (red). An exponential curve is then fitted on measured ablation velocities, providing an initial recoil velocity as estimation for the cortical tension. C: Ablation speeds differ between two sides of the polarizing zygote, and between two directions of ABpl. 95% confidence intervals and p-values were approximated using bootstrap, 2000 samples. The difference is significant with p < 0.0005 for both cases. D: Comparison of cortical tension measurements with tangent FFI tension inference for the 7-cell stage embryo. The positive correlation is significant (p < 0.01, using bootstrap, 10,000 samples). E: Comparison of cortical tension measurements with FIDES tension inference for the 7-cell stage embryo. The positive correlation is significant (p < 0.0001, using bootstrap, 10,000 samples). F: Exponential decay rate for ablations of the 7-cell stage cells, showing 95% CI.

As cortical laser ablation requires a complex and precise experimental protocol, we first validated our procedure on the polarizing zygote (P0). At this stage, the anterior side has more actin and myosin, and contains highly contractile bundles that ruffle the membrane. Mayer et al. (2010) used cortical laser ablations to measure surface tensions at this stage, and the anterior side was found to have roughly twice the tension of the posterior. Following the experiments of Mayer et al. (2010), we made 8 µm cuts parallel to the AP axis at the 70% retraction stage. We found average initial recoil velocities on the anterior side (P0-A) of 18.7 µm/s [16.4–20.9 95% CI], and 10.2 µm/s [8.2–12.5 95% CI] on P0-P, the posterior side (see fig. 3C, see Supplementary Table S1 for full results). This difference was highly significant, and matched the roughly two-fold tension difference reported by Mayer et al. (2010).

Next, we performed ablations on the 7-cell stage *C. elegans* embryo to sample the surface tensions. For this stage the cuts could only be 5 µm long, due to the reduced cell size. We initially assumed surface tension to be relatively homogenous and isotropic in this stage, and cuts were made parallel to the anterior-posterior (AP) axis. However, while performing the experiment, we noticed that specifically for the ABpl cell there was strong anisotropy in tension depending on the direction of the cut, with the AP cut (fig. 3C, green arrow) having much higher initial velocity than the perpendicular dorsal-ventral (DV) directed cut (fig. 3C, purple arrow). When cut parallel to the AP axis, ABpl had an average velocity of 13.5 µm/s, significantly higher than the 7.2 µm/s when cut perpendicularly (p < 0.0005). This shows that ABpl has highly anisotropic tension in this stage, with a higher tension in the DV axis than in the AP axis. The remaining cells at this stage cells had a similar response in both directions. For comparisons between cells, we use the average of both directions as a measure of the total surface tension in ABpl, following the derivations in Fratzl et al. (2021). All ablation data is included in the Zenodo package.

Next, we compared the experimental measurements of the 7-cell surface tensions, to those inferred by both FIDES and tangent FFI. We applied both methods on data from 11 wild-type *C. elegans* embryos. Bootstrapping was used to assess significance of trends and provided 95% confidence intervals for inferences and measurements. Figure 3D shows that tangent FFI has a modest correlation with the ablation results, with a Pearson’s r of 0.45 (p < 0.01). By contrast, FIDES has a better correlation, with a Pearson’s r of 0.69 (fig. 3E, p < 0.0001). Curved FFI performed considerable worse (see Supplementary Figure S4C).

When comparing cells (Supplementary Figure S5), we observed that cells P2 and E have the highest ablation velocities measured at the 7-cell stage, while also having the fastest exponential decay (fig. 3F). This implies that both have cortices with high tension and high stiffness (see Materials and methods). The high values for P2 can be explained, as it is performing mitotic rounding in preparation for cell division, which increases both surface tension and stiffness (Taubenberger et al., 2020). However, the non-dividing E cell stands out due to its relatively high stiffness and apparent higher surface tension than what was predicted based on its shape (fig. 3E).

### Force inference for early embryogenesis and comparison to earlier work

With FIDES, we analyzed 15 wild-type embryo time-lapses up to the 8-cell stage, comprising a total of 210 time points (see Supplementary Figure S6). Figure 4 shows our aggregated surface tension and adhesive tension inferences for several snapshots between the 2- and 8-cell stage. The surface tension distribution is highly dynamic, both between different cells and over time. The most prominent factor is mitotic rounding, where cells increase their surface tension to round up in preparation for division. In stage 4A, cells all have similar surface tension, while the later stage 4B features two cells primed for division. In Supplementary Figure S7, we show a smoothed tension profile for every cell, continuous over time. These profiles indicate surface tension differences between cells rather than absolute values, as the inferred tensions are normalized making the reported cell surface tensions average 1. The corresponding evolution of cell pressure can be seen in Supplementary Figure S8.

**Fig 4.**
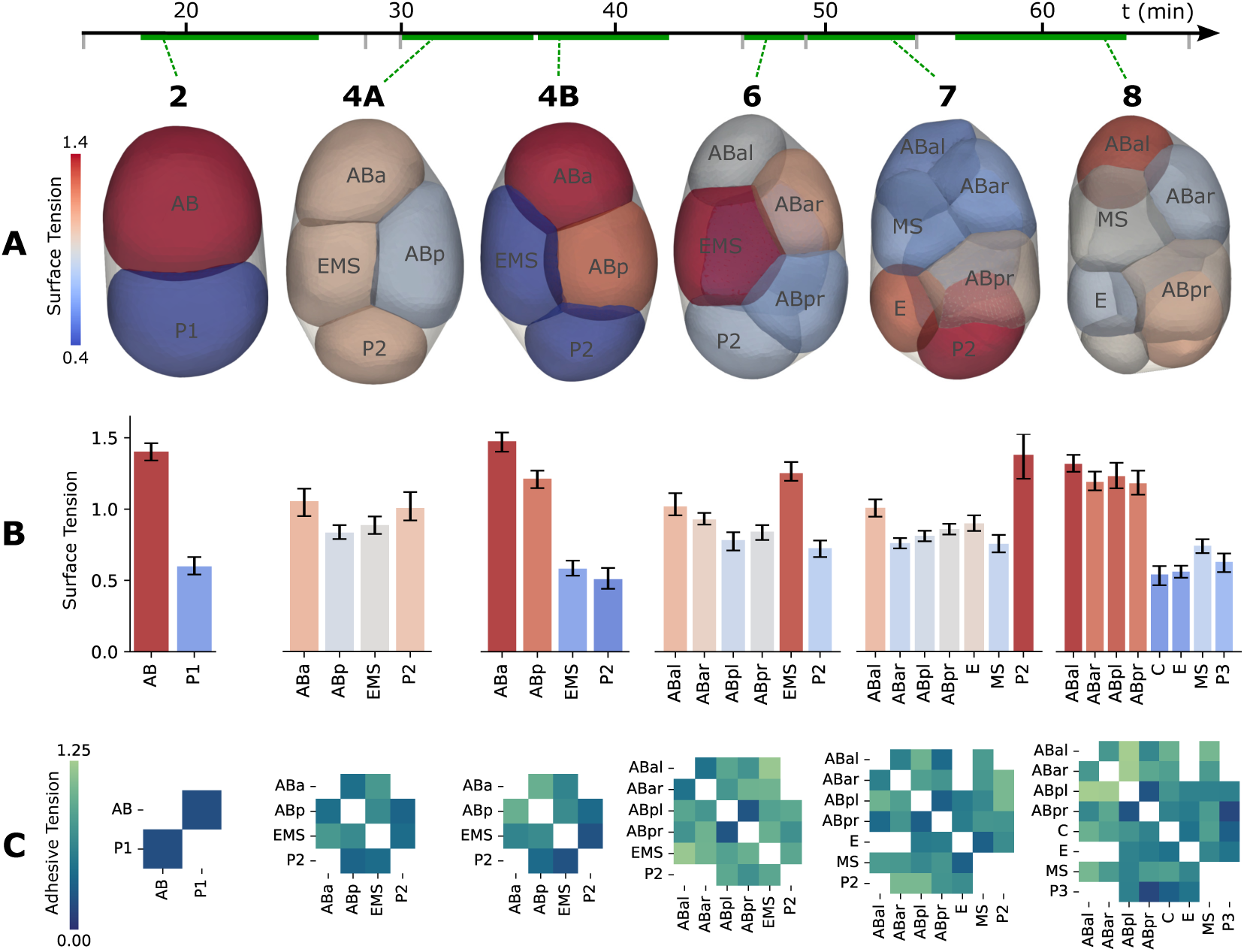
FIDES tension inference over early development. A: We show six snapshots of a developing *C. elegans* embryo, each representing the aggregated force inferences at their respective cell stage. The 4-cell stage is split up into two columns (4A and 4B), as it is a long-lasting stage with significant ongoing changes. B: Inferred cell surface tension results for every cell. The surface tension bars have 95% confidence intervals, calculated via bootstrap, 1000 samples. C: Inferred adhesive tension results for every cell-cell contact.

Adhesive tensions are contact-specific and visualized as heatmaps in fig. 4C. The average inferred adhesive tensions increase over time, gaining more importance relative to cell surface tension. FIDES infers minimal adhesive strength for the single contact at the 2-cell stage, indicating that the cell shapes can be primarily explained by surface tension differences between AB and P1 and confinement of the eggshell. As cell stages progress, stronger adhesive contacts appear.

For comparison, we also performed FFI on the same embryo time-lapses. Supplementary Figure S9 shows the correlation between FIDES and tangent FFI for every frame, with the correlation coefficient gradually decreasing. The two methods are in agreement for the very passive 2- and 4-cell stages, but start to diverge from the 6-cell stage onward. This mirrors the results from the synthetic embryo, where FIDES outperforms FFI when there is confinement and anisotropic force generation.

Next, we compare our inferences to previously published analyses on the *C. elegans* embryo. In a study by Yamamoto and Kimura (2017), the eggshell was removed from the 4-cell stage embryo, to make inferences on cell attraction using a model for interacting spheres. They found that the adhesion strength between EMS and P2 was significantly lower than the contacts involving ABa, ABp, and EMS. As shown in fig. 5A, this matches our inference with FIDES. Using bootstrapping, we confirm that the adhesive tension between EMS and P2 is significantly lower than the other contacts measured by Yamamoto and Kimura (2017) (p < 0.0005). Additionally, we see that this is also the case for the adhesive tension between ABp and P2, which was not previously observed as the cells were physically separated with the eggshell removed.

**Fig 5.**
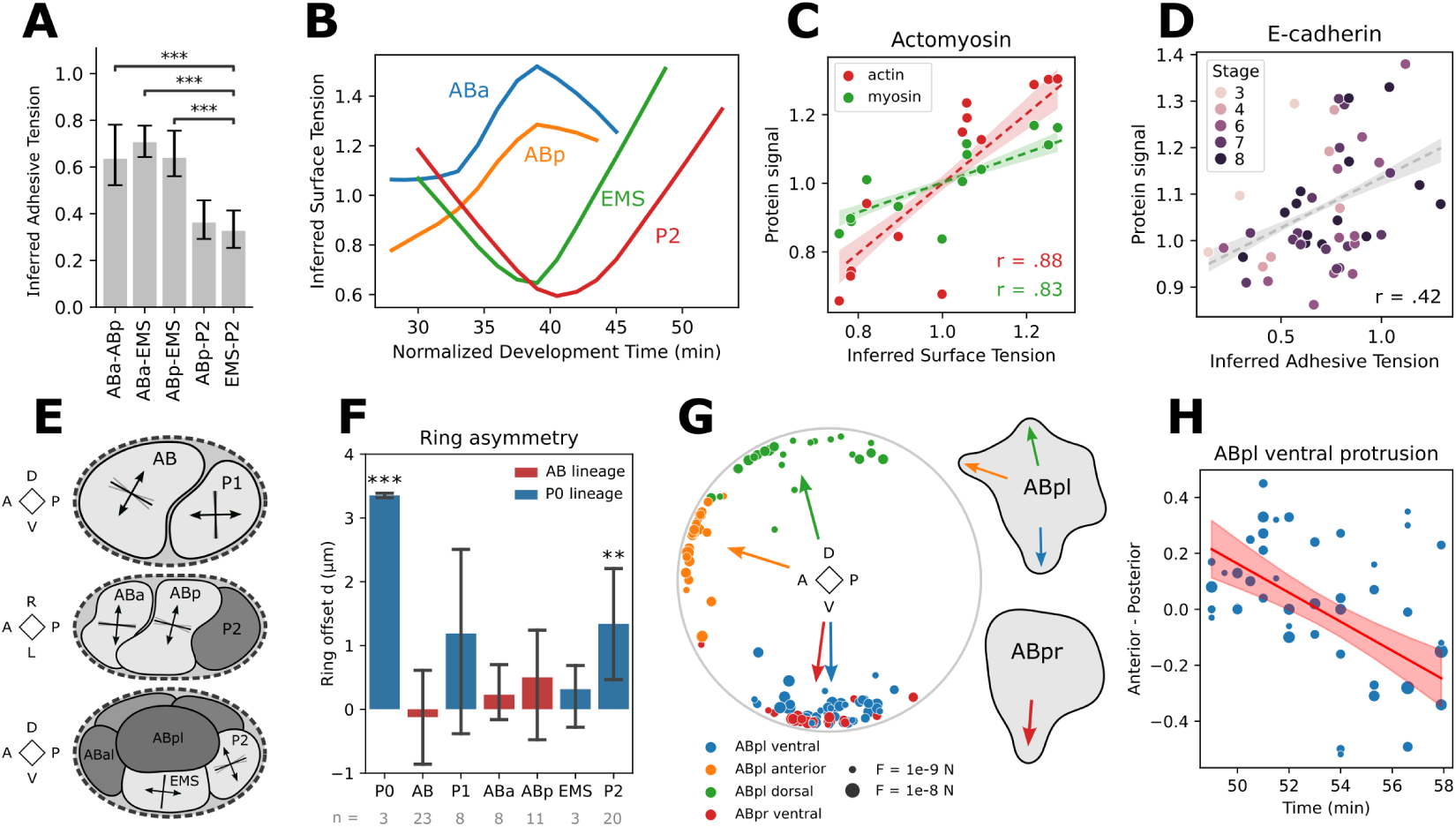
Inference comparisons to literature and protein presence. Data is aggregated from 15 time-lapses (see Supplementary Figure S6). A: Inferred adhesive tensions of the 4-cell stage (n=43). Bootstrap (2000 samples) was used to calculate 95% confidence intervals. B: Timeline of inferred surface tension for the 4-cell stage cells. Data is smoothed using a lowess regression. Surface tensions are normalized so they average 1 at each time instance. C: Comparison of inferred surface tension to experimental measurements of cortical actin and myosin, measured across the free cell surface. Points are aggregated per cell for all cells between the 2- and 8-cell stage. Regression band with 95% confidence interval via bootstrap, 1000 samples. D: Comparison of inferred adhesive tension to experimental measurements of E-cadherin on the cell-cell interfaces, aggregated per contact for different cell stages. Regression band with 95% confidence interval via bootstrap, 1000 samples. E: Orientation of inferred cytokinetic rings for different cell stages. Division planes are projected to the most meaningful axis and standard deviations of the planes are shown in gray. F: Asymmetry of inferred cytokinetic ring offsets for every cell division, with 95% confidence intervals. Difference from zero is significant for P0 (p=0.00004) and P2 (p=0.009). G: Orientation of inferred protrusions for ABpl and ABpr. Individual observations are 3D unit vectors projected on the plane perpendicular to the LR axis, while the weighted average orientation is shown as an arrow. H: Rotation of the ventral protrusion of ABpl over time. Weighted least squares regression finds a significant effect of the AP axis component over time (p<0.0001), meaning that the protrusion moves from pointing slightly posterior to slightly anterior over the course of 10 minutes.

A follow-up experiment was performed by Yamamoto et al. (2023), where they used Atomic Force Microscopy to measure the surface tension at the 2- and 4-cell stages. They found that AB had significantly higher tension than P1, and the daughters of AB had higher tension than the daughters of P1. This is a difference that arises during the asymmetric cell division of the zygote P0. During the first division, the AB cell receives a greater share of cortical actin and myosin compared to its sister P1, and this asymmetry persists through multiple generations (Caroti et al., 2021). We also observe that AB and its descendants generally exhibit higher surface tension than the other cells, as seen in stages 2, 4B, and 8, but notably not in the 7 cell stage (fig. 4B).

Cortical tension varies during the cell cycle, and typically reaches its highest point during mitotic rounding (Taubenberger et al., 2020). Up to metaphase, a cell’s tension, stiffness, and pressure all increase, only to drop at anaphase, as the cell starts elongating before division. Figure 5B shows the measured surface tension of the 4-cell stage cells. The anterior descendants (ABa, ABp) exhibit a high average tension and clearly show mitotic rounding in their profile. In contrast, the posterior cells (P2, EMS) also build up some surface tension towards the end, but their profile decreases first. It is important to note that these inferences are relative, and trends over time may be influenced by changes in the properties of other cells. We discuss these profiles further in the Discussion. Supplementary Figure S10 gives a detailed view on all cell surface tensions leading up to division, with the individual measurements.

Surface tension and adhesive tensions are generated by cytoskeletal components that can be directly imaged. We obtained confocal fluorescence microscopy data to quantify fluorescent markers for actin, myosin, and E-cadherin during early embryogenesis. This data was normalized per frame, to average 1, so we can make a fair comparison to the relative values from inferred surface tension. Figure 5C shows that the surface tension of every cell, averaged over their lifetime, correlates well with both their actin (r=0.88, p=0.0001) and myosin (r=0.83, p=0.0005) quantities on the free surface. These results also highlight the previously mentioned effect that asymmetry in cortical F-actin and myosin arising in the zygote, higher in AB and descendants, correlates with a higher tension. Figure 5D show that the cell-cell adhesive tension during different stages has a positive correlation with their E-cadherin signal on the interface (r=0.42, p=0.003).

In fig. 5E, we analyze the orientation of cytokinetic rings positioned by FIDES. During the two-cell stage, AB’s division causes the plane to rotate away from the AP axis. P1 is subsequently pushed to one side of the eggshell, and divides along the AP axis. These results are consistent with the final spindle positions in AB and P1, as shown by Pimpale et al. (2020). During the four-cell stage, we see the posterior shift of future ABar and ABpr as they divide, with the division planes slightly tilted.

This asymmetry is caused by a skew of the spindle in ABa and ABp, and later defines the left-right (LR) axis (Pohl and Bao, 2010). After that, EMS divides along the AP axis, and P2 divides along the DV axis, slightly tilted, with future P3 towards the posterior. The ring orientations of all observed cells align with previous observations of cell positioning (Rose and Gonczy, 2014). Full results for the orientation and total force of the rings and protrusions are provided in Supplementary Table S2.

FIDES is able to model asymmetric divisions, giving the division plane an offset distance from the cell center. Figure 5F shows the summarized offset data of all divisions. We found that cells P0 and P2 have a significant offset (p-values 0.00004 and 0.009, respectively). This is consistent with the literature, as the P lineage has the strongest asymmetric cell divisions (Fickentscher and Weiss, 2017). The average asymmetry in P1 does not reach significance due to high variability, and would need more observations.

During the 7-cell stage, we modeled protrusions for ABpl and ABpr. Figure 5G illustrates the orientation of the inferred protrusions in every sample, with the weighted average indicated as arrows. We reliably find the ventral protrusions of ABpl and ABpr, as well as ABpl’s dorsal lamellipodium and one of the anterior protrusions, as described by Pohl and Bao (2010). We further observed that ABpl’s ventral protrusion changes orientation over time (fig. 5H). A weighted least squares fit reveals a significant shift from posterior to anterior (p < 0.0001). This movement ends with the ventral protrusion being positioned on MS, with E-cadherin HMR-1 on their interface (Pohl and Bao, 2010; Caroti et al., 2021).

## Discussion

We presented FIDES, a new method for cellular force inference that uses mechanical simulation of cell shapes derived from segmentation results. The simulation framework can account for common features of developing systems, including spatial confinement, and local force generation related to cytokinesis and protrusion formation. FIDES fits parameters to best explain cell shapes, including the surface tension for every cell, adhesive tension for every existing cell-cell contact, and various parameters defining the models for local forces. We evaluated FIDES performance and compared it to the current state-of-the-art FFI. We first validated our method on synthetic embryos, and found that it can predict the surface and adhesive tensions accurately, even in the presence of cell division and protrusion formation. FFI performed equally well on simple foam-like structures, but had a higher error in more complex cases where local forces and eggshell confinement distorted the cell shapes. Curved FFI performed relatively poorly, as it partly relies on membrane curvatures, that are severely impaired by cell divisions and confinement. Meanwhile, tangent FFI is robust against distortion of curvature, but its performance can still suffer due to unreliable junction angles when cells press against the eggshell (table 1). It should be noted that FFI methods did have a positive correlation with the solution in our synthetic examples. They may further be applied together with FIDES analyses, for instance as an initial guess of tensions to speed up the optimization. We next performed experimental validation for surface tension predictions at the 7-cell stage of the *C. elegans* embryo, using cortical laser ablations.

FIDES and tangent FFI both had a positive correlation, but FIDES yielded better results. FFI methods are apparently less well-suited for multicellular systems that don’t resemble a passive, quasi-static foam, whereas the FIDES model can process more complex cases. Further, FIDES evaluates the entire cell shape and, in contrast to FFI, is not solely reliant on local features such as contact angles which are sensitive to segmentation errors, especially for 3D segmentations.

During our ablation experiments, we found that the ABpl cell had a highly anisotropic surface tension, with a higher tension in the direction of the DV axis than the AP axis. We attribute this to the ventral protrusion of ABpl, which forms at the 7-cell stage and initiates a significant movement in the compressed embryo (Pohl and Bao, 2010; Jelier et al., 2016). Given that ABpl moves ventrally, this appears a reasonable explanation for the additional surface tension in the direction of the DV axis. An unexpected result was the ablation profile of cell E, which implied a higher tension than expected based on the prediction by FIDES, and higher stiffness than most other cells at this stage. It appears that the cortex of E is different from other cells, in that it exhibits high tension and stiffness, while simultaneously showing a lower non-muscular myosin and F-actin signal. Further, as we previously reported, the E cell has a cortex structure that is much more dynamic than its sister cell MS, as well as other cells at this stage, and arose following a Wnt-signaling-induced asymmetrical division (Caroti et al., 2021). Differences in thickness of the cortex, as well as other biomechanical changes such as lower cortex viscosity, are not captured in the simple model we used here for interpreting the laser ablations. More advanced models have been proposed that better use the information in images of cortical ablations (Saha et al., 2016), but no bioinformatics toolbox is available to apply these, so we leave this for future work.

We used FIDES to analyze wild-type *C. elegans* embryos between the 1- and 8-cell stage, and inferred forces over time. We see strong effects of the cell cycle, as mitotic rounding strongly increases cortical tension during the metaphase, followed by a decrease before division. This effect is clear for the anterior cells, yet distorted for the posterior ones fig. 5B. Given that tension inferences are normalized, surface tensions may appear to decrease simply because others are increasing more. We argue that this is the case for the posterior cells as they have lower overall surface tension than their anterior neighbors, which are dividing at the same time. Further, surface tension tends to vary by lineage, where the descendants of AB tend to have a higher tension than those of P1, except in the 7-cell stage as noted. This trend in cell-averaged surface tension correlates strongly with actin and myosin measurements, implying that, in general, in the early *C. elegans* embryo more cortical material is associated with a higher tension. We find a weaker, yet positive correlation between contact-specific adhesion energy and E-cadherin measurements, confirming the importance of the adhesion protein during early *C. elegans* development. Maître et al. (2012) found similar results in zebrafish, where E-cadherin decreases the cortical tension in the interface, as well as physically coupling cells together, which together increase the cell contact area. Parametrization of local force models by FIDES could verify previously reported observations of division orientations, asymmetry, and protrusion formation. We further observed a posterior to anterior shift of ABpl’s ventral protrusion, which matches previous observations (Pohl and Bao, 2010; Caroti et al., 2021).

The FIDES method has some limitations that provide opportunities for further work. First, the FIDES shape models do not fully explain the observed shapes, but instead are aimed to capture the largest effects. The models can likely be improved and expanded as they are used more broadly. New active force models would need to be added to FIDES to capture other events that happen later in *C. elegans* embryogenesis, or in other organisms. Second, the models will need to be further validated. In this paper we performed extensive, intricate laser ablation experiments to confirm surface tension predictions of the 7-cell stage. Other inferences were compared to previous reports on surface tensions, adhesive tension between cells, division angles, and protrusion formation. However, to ascertain the accuracy and limits of the FIDES models, for example of the adhesive tension inferences, would require further quantitative experimental validation. It is also relevant to note that our simulations require several physical parameters to be set (see Supplementary Table S3), many of which have a degree of certainty for *C. elegans* embryos and other systems. Most of these parameters, however, describe friction and viscosity, primarily influencing the speed of reaching static equilibrium shapes in our simulations rather than the shapes themselves. Third, similar to other methods for image-based force inference, FIDES provides relative, and not absolute, force estimates. An experimental measurement is needed to estimate the absolute values at different stages. Without such grounding of the inferences, we can compare the relative tensions within a cell stage, but it is more difficult to compare tensions over time. Fourth, FIDES analyzes a single time point to infer forces, making it unable to capture dynamic cellular movements. While we make the assumption that the cell shapes are in static equilibrium, a developing embryo likely experiences viscous forces that may affect cell shape. In an earlier study regarding cell division in *C. elegans*, Cuvelier et al. (2023) showed that cortical tension forces dominate over viscous forces, limiting the error introduced from ignoring viscous forces during dynamics. Future work would extend this method to analyze cell shape changes in time-lapses, so that dynamic forces may be inferred more accurately. This would allow us, for example, to investigate how the ABpl protrusion helps the rotation of the 7-cell *C. elegans* stage, and other cell movements such as gastrulation. Lastly, FIDES is computationally intensive compared to the efficient analytical methods employed by FFI methods, which is a consideration when large numbers of cells are involved and the assumptions of quasi-static, passive foam-like cellular behavior mostly hold. More efficient formulations and optimization approaches can also be explored to make FIDES converge to a solution faster.

Image-based force inference approaches hold the promise to readily provide researchers with accurate data on cell generated forces. With our experimental model, we tracked tensions over time in the *C. elegans* embryo and also identified local force generation. Our results reveal a remarkably dynamic system with substantial surface tension differences between cell lineages, distinct behavior of individual cells, and rapid changes in the tension distribution over time. Characterizing force generation during development yields valuable information, in particular to understand the different active contributions involved in cellular positioning. Broad application of image-based force inference can facilitate the study of various cellular processes in animal systems, including morphogenesis and regeneration. While FFI methods are fast and useful in specific scenarios, we demonstrated limitations that constrain their applicability. In this paper, we illustrated that by employing a flexible biophysical modeling system, FIDES can go beyond these limitations and make accurate inferences in a system with active movements.

## Materials and methods

### *C. elegans* Strains and maintenance

*C. elegans* strains were grown on NGM plates at 20℃ as described by Brenner (1974). For cell segmentations we used RJ013, a cross between LP306 (cpIs53 [mex-5p::GFP-C1::PLC(delta)-PH::tbb-2 3’UTR + unc-119 (+)] Il (Heppert et al., 2016)) and SWG001 (gesls001 Pmex-5::Lifeact:mKate2::nmy-2UTR, unc-119+1(Reymann et al., 2016)). For ablations, we used RJ012, a cross of LP162 (nmy-2(cp13(nmy-2::GFP + LoxP)) I. (Dickinson et al., 2015)) and SWG001. Both strains were previously documented by Caroti et al. (2021).

### Imaging and embryo segmentation

Live embryos were mounted on slides with M9 buffer and compressed to 20 µm with Polybead Microspheres (Polysciences). When working with zygotes, 4% sucrose was added to the medium to stabilize the membranes (Mayer et al., 2010). All imaging was done using a Zeiss LSM 880 confocal microscope with a Plan-Apochromat 63x/1.4 DIC M27 oil immersion objective. We gathered image stacks every 90s, with a z-step of 1 µm. 3D cell segmentations from *C. elegans* embryos were obtained with the spheresDT/Mpacts-PiCS procedure (Thiels et al., 2021). To make the cell meshes resulting from the segmentations more suitable for further simulation procedures, we optimized the simulation relaxation step. Specifically, a convex hull of the cell nodes is computed during the relaxation, and added as an approximation of the eggshell. The addition of an eggshell is helpful as a boundary condition for the segmentation pipeline, but is also crucial for the simulations, as it serves as a rigid confinement ad push-off point for the cells. The mechanical parameters of the DCM were also modified, featuring lower viscosity to guarantee proper cell contacts (Supplementary Table S3). After relaxation, we manually aligned the embryos to a common coordinate system (Supplementary Table S2). The processed cell meshes are then used by both FIDES and FFI.

For quantitative measurements of actin, myosin, and E-cadherin, we used the same microscopy setup as described above. Processing of the fluorescent signal is described in Supplementary Text 1.2.

### Cell mechanical model

All embryo simulations are performed using Mpacts, a particle-based simulation framework using the Discrete Element Method to compute the motion of interacting particles based on a force balance (Odenthal et al., 2013; Smeets et al., 2019). Cells are modeled as a foam-like material using the Deformable Cell Model (DCM), which considers the cell cortex and membrane as a viscous shell under tension. Forces like tension and contact pressure act on the nodes of a triangular mesh, causing realistic deformation of the cell shapes. The cells are parametrized by cortex thickness *t*, surface tension *γ*, and viscosity *η*, as shown in fig. 1A. They have a reference volume *V^∗^*, and experience resistance to volume change controlled by a bulk modulus *K*, which exerts additional pressure *P* on the cell membrane. Cell-cell interactions are managed through repulsion and contact-specific adhesive tension *ω*. When cells are isolated and free from additional force models, surface and adhesive tension together determine the contact angles at triple junctions, in accordance with the Young-Dupré equation. Validations of the DCM regarding geometry and contact angles are provided in Cuvelier et al. (2023). At the cell-cell interfaces there is also wet friction with friction coefficient *ξ*, while medium viscosity acts on all nodes. A more in-depth description with a force balance per node is included in Supplementary Text 1.1.

### Contractile ring model

Preceding cell division, F-actin fibers align and converge into a contractile ring, which gives rise to the cleavage furrow (Eweis and Plastino, 2020). The spindle location guides the positioning of this ring, and any spindle offset from the cell center will result in an asymmetric cell division. We approximate this process by adding additional local forces on the existing DCM, similar to Cuvelier et al. (2023). The ring is defined by four parameters: two spherical coordinates (*θ*, *ϕ*) for the normal direction of the division plane, a spring constant *k*, and an offset distance *d* (fig. 1C). Along the intersection of the division plane and the cell mesh, a ring of nodes is selected to become the furrow ring. These nodes are then connected using linear-elastic springs, with spring constant *k*, resulting in inwards force *F_R_*. To achieve a smooth furrow, *F_R_* is distributed over the area around the division plane, preventing a sharp crease. Asymmetric division is achieved by altering *d*, which positions the division plane at an offset distance from the center of the cell. However, this setup results in two sides of the cell having the same surface tension, but different curvatures, which is an unstable configuration (Sedzinski et al., 2011). To prevent one of the sides collapsing, we add proportional controller *K_D_*, which resists changes in cell area on both sides of the division plane, resulting in pressure forces *F*_*P*1_ and *F*_*P*2_, similar to Cuvelier et al. (2023). The implementation of the contractile ring model is further explained in Supplementary Text 1.1.

### Protrusion model

The protrusion model has four parameters, two spherical coordinates for the direction, a total protrusive force *F_P_*, and an angle *α*. As illustrated in fig. 1D, the spherical coordinates are projected from the cell’s center to establish a projection axis, which serves as the central axis of the protrusion. Total force *F_P_* is then distributed using a Difference of Gaussians *G_a_* − *G*_2*a*_, projected perpendicular to the membrane. This gives the center of the protrusion a net outward force, while the edges are pulled inwards to support the protrusive action. To guarantee that the net force is zero, the remaining sum of all individual forces is inverted and redistributed over the protrusion. The implementation of the protrusion model is further explained in Supplementary Text 1.1.

### FIDES Optimization

The FIDES procedure performs error minimization by repeatedly evaluating brief Mpacts simulations on the embryo shapes from segmentation, while varying the following mechanical parameters:

- Surface tension per cell (*γ_i_*).
- Equilibrium volume per cell *V_β_*\*.
- Adhesive tension per cell-cell contact (*ω_ij_*).
- Four parameters defining each protrusion (*α*, *F_p_*, *θ*, *ϕ*).
- Four parameters defining each contractile ring (*d*, *k*, *θ*, *ϕ*).

Each simulation lets the cell shapes deform under the DCM for dimensionless time *ηt_c_/γ* = 2.5, where *η* is cortex viscosity, *t_c_* is cortex thickness, and *γ* is the average surface tension. At the end of a simulation, we compute the deformation error *E_D_*, as the sum of all raycast distances *d_i,α_* = *||***x_i_** *−* **x***_α_||*, where **x_i_** is the initial position of node *i* and **x***_α_* is the intersection point between the ray from point **x_i_** and node normal **n̂***_i_* with the deformed mesh at *t* = 2.5*ηt_c_/γ*. Such that *E_D_* = ∑_i_ *A_i_d_i,α_* with *A_i_* the voronoi area of node *i*. In addition to the deformation error, we add a linear penalty term for total adhesion energy, contractile ring force, and protrusion force, so these are only kept if they significantly benefit the solution.

The optimization process is shown in Supplementary Figure S11. We start with an initial proposal, which assigns every cell equal surface tension and counteracting pressure to maintain its volume, while every unique cell-cell contact gets equal adhesive tension. The average cell surface tension is fixed, as there is no reference for the real surface tension over time. Depending on the cell stage, local force models are added to the simulation, each with its own set of parameters. These local force models include a contractile ring for cells approaching division, and a protrusion model for cells like ABpl and ABpr during the 7-cell stage. In vivo, ABpl is observed to have a dorsal lamellipodium, a ventral protrusion, and anterior filopodia, while ABpr has a ventral protrusion (Pohl and Bao, 2010). We view each of these as a simple protrusion, only differing in their initial orientation. The initial orientation of rings and protrusions is defined by a manually estimated schedule, based on a synchronized time-lapse. The embryo is manually rotated to align with the expected orientation, and the frames are given a cell stage timestamp, which make it possible to automatically initialize the rings and protrusions and have them find their right place and properties throughout the optimization process. Given the invariant development of *C. elegans*, this fixed placement of rings and protrusions is an effective solution. However, we also investigated the possibility to automatically detect these using the FlowShape (van Bavel et al., 2023). This method is quite accurate at both the detection of protrusions and cytokinetic rings, with an accuracy of 90% and 97% respectively. Further details are described in Supplementary Text 1.3.

During the FIDES procedure, the parameters undergo stochastic optimization (Spall, 2011) with up to 150 iterations. At each iteration, each parameter is randomly perturbed using a Gaussian distribution, followed by a separate simulation to check if the deformation error improves compared to the baseline of the iteration. At the end of an iteration, all parameter changes that had a positive effect are applied to the baseline solution, to be used for the next iteration. The optimization process stops when the error does not decrease for 15 successive iterations, or the iteration limit is reached. Note that, gradient descent was tested as an alternative optimizer, converging to a solution slightly faster than stochastic optimization, but occasionally failing due to the non-smooth error landscape. Given good quality meshes, both methods yield similar and consistent results, despite the random effects from stochastic optimization. Reported surface and adhesive tensions are always normalized as *γ/γ*_0_ and *ω/γ*_0_, with *γ*_0_ the initial surface tension, making the reported surface tensions average 1.

The computational time is highly dependent on the number of cells, as more cells increase both total triangle count and the number of parameters. Mesh density could be lowered to speed up the simulations, but we kept it set to the recommended density from the segmentation procedure to maintain all shape information. We ran the optimizations on an 18-core Intel Xeon Gold 6140 CPU, with a base clock of 2.3 GHz. Every iteration is parallelized, and a single simulation thread took around 10 seconds for the 2-cell stage. The total computational time varies, with 2-cell stage embryos taking around 10 minutes and 7-cell stage embryos requiring approximately 2 hours. In total, 210 time steps from 15 different wild-type embryos, up to the 8-cell stage were processed.

### Foam force inference

Curved FFI is performed based on the algorithm of Xu et al. (2018). One major difference is that we start from triangular meshes and contact data, which we obtained via spheresDT/Mpacts-PiCS. Where three cells meet, or two cells and the exterior, a triple junction is defined (fig. 1E). To measure the angles of a triple junction, the triangles from the cell meshes are first reassigned to three distinct surfaces. Where two cells make contact, we consider the two contacting surfaces as one interface a shared interfacial tension. At several points along the junction, we then measure the angle between the surfaces in a plane normal to the contact line. The triple junction angles are averaged along the junction to give us the three final angles *θ_ij_*. We omit measurements if the contact surface was less than 50 triangles, as this is considered too small for robust measurement.

For surface curvature measurements, we consider every surface not part of, or directly adjacent to, a triple junction. By averaging the principle curvatures along a surface, we arrive at one mean curvature per surface. The curvature at cell-cell interfaces is defined to be the average curvature of both membranes. Also, curvature measurements are omitted if the contact area contains fewer than 50 triangles, as we consider this too small to be reliable.

Junction angles *θ* and mean curvatures *H* allow us to retrieve information about pressure *P*, surface tension *γ_i_*, and interfacial tension *γ_ij_*, or alternatively, adhesive tension *ω_ij_*. The following equations likewise consider the interface between cells with an interfacial tension *γ_ij_* = *γ_i_* + *γ_j_− ω_ij_*. The first equation used is the Young-Laplace equation, relating a pressure difference to the mean curvature and tension of a surface.

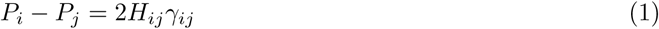

Second, we use the Young-Dupré equation to relate the contact angles to the tensions of the three surfaces.

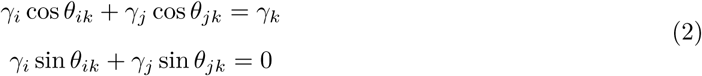

We get one equation for every curvature, and two equations for every set of angles from a triple junction. The unknowns are one pressure and surface tension per cell, and one interfacial tension per cell-cell contact. We add another equation to remove the scaling ambiguity, forcing the average surface tension to be 1. From the 2-cell stage onward, this produces an over-determined system of equations, which we solve using the pseudo-inverse matrix.

Tangent FFI is performed in a similar way, but ignores surface curvature and does not use Young-Laplace equations. Consequently, no pressures are obtained from solving the resulting system either.

### Cortical laser ablation

Cortex ablations were made using a 355 nm 1 kHz pulsed laser UGA-42 Caliburn from Rapp OptoElectronic. This system was mounted on the Zeiss LSM 880 confocal microscope used for imaging, but we now image one z-slice at *±*1 frame per second. Nmy-2::GFP and Lifeact::mKate2 were excited using 488 nm and 561 nm lasers respectively, and imaged using the Airyscan detector in R/S mode. After focusing on the cell cortex, the laser was activated for 500ms to rupture the cortex over a 5 µm line (8 µm for zygote cuts). Measurements were only considered valid if the surrounding material showed a response to the cut, followed by repair of the cortex, which excludes irreversible ruptures. Zygote cuts were made parallel to the AP axis, when P0 was at the 70% retraction stage, as described in Mayer et al. (2010). For the 7-cell stage cells, we timed the cut to be in between EMS and P2 division, a time interval of only a few minutes.

A crucial part of the procedure is the calibration of the ablation laser set-up. The applied laser power needs to be carefully adjusted, as too little power will not reliably ablate the cortex, while too much power will kill the cell and induce irreversible membrane rupture. Additionally, the focus depth of the ablation laser was determined to be offset by 1 µm relative to the imaging plane. To compensate, the z-position was briefly adjusted to the optimal depth during the 500ms cutting interval.

To quantify the recoil movement after ablation, we used a custom Particle Image Velocimetry (PIV) implementation, based on manually tracking cortical highlights and resulting in 9000 markers. This solution was chosen as automated PIV algorithms had trouble with high levels of noise in the fluorescent images. Cortical tension and stiffness were then inferred from the recoil movement by modeling the actomyosin cortex as a Kelvin-Voigt material, including a spring, dashpot, and an active contractile element as in Mayer et al. (2010). The underlying assumption is that, on short timescales, the actomyosin cortex behaves as an active viscoelastic material. Given the mechanical tension *T*, active contractile tension *C*, elastic stiffness *k*, and damping coefficient *ζ*, we define the displacement during ablation as:

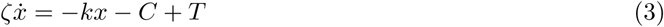

Before ablation, tension *T* and contractility *C* are in balance, resulting in an equilibrium. When the cortex is ablated, the mechanical tension *T* becomes zero, and the cortex will recoil with velocity:

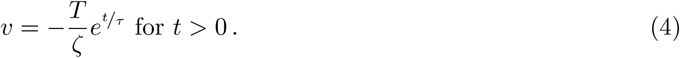

Where relaxation time *τ* = *ζ/k*. This means that the cortex will recoil away from the ablation line, following an exponentially decaying profile. Assuming constant damping, the cortical tension is proportional to the initial recoil velocity after the ablation, while cortex stiffness is inversely proportional to the relaxation time.

Figure 3B shows an exponential curve fitted to the tracked velocities using non-linear least squares. However, since every measured velocity is itself an average of the real velocity over a time interval of about one second, we end up underestimating the initial velocity. This effect becomes more pronounced when the relaxation time is short, as shown in Supplementary Figure S12A. After simulating an ideal ablation and seeing the effect of a one-second averaging window (Supplementary Figure S12B), we found that we can correct the initial velocity using the measured exponential decay.

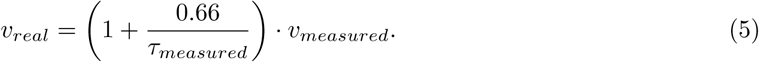

Each experiment has many tracked markers, following the deformation of the cortex. To obtain confidence intervals, all individually tracked points were grouped by condition, and weighted bootstrapping was performed (2000 samples). The weights ensure that different cell cuts had an equal contribution, regardless of the number of sampled points per cut. This results in 2000 exponential curve fits per condition, whose parameters are then used to get the average and 95% confidence intervals. When a total surface tension for ABpl was required, we took the average of both directions. P-values were simulated using the same bootstrap samples, considering the fraction of times one condition scored higher than the other.

## Acknowledgments

We thank Stephan Grill for helping us set up the ablation protocol. Additionally, thanks to Jiří Pešek for help with Docker. Experiment expenses were supported by FWO Research grant number G008423N. M.V., W.T., J.V., and C.v.B. were supported by FWO Aspirant grants 1194222N, 11I2921N, 11D9923N, and 11L0923N respectively.

## Supplementary Information description

### Supplementary Text

**Text 1:** DCM force balance

**Text 2:** Protein signal correction

**Text 3:** Automatic detection of protrusions and cytokinetic rings

### Supplementary Figures

**Figure S1:** Interfacial and adhesive tension at the cell-cell interface

**Figure S2:** Protrusion model

**Figure S3:** Sensitivity analysis for FIDES physical parameters

**Figure S4:** Synthetic and ablation performance for curved FFI

**Figure S5:** Synthetic and ablation performance for curved FFI

**Figure S6:** Embryo time-lapses and their coverage of early development

**Figure S7:** Timeline of inferred FIDES surface tensions between 1- and 8-cell stage

**Figure S8:** Timeline of inferred FIDES pressures between 1- and 8-cell stage

**Figure S9:** Correlation between results from FIDES and tangent FFI

**Figure S10:** Surface tension inferences during mitotic rounding

**Figure S11:** Parameter optimization strategy for FIDES.

**Figure S12:** Correction for laser ablation velocity

### Supplementary Tables

**Table S1:** Numeric ablation results and sample sizes

**Table S2:** Averaged parameters of local force generating activities

**Table S3:** Mpacts physical parameters

### Supplementary Videos

**Video 1–3:** *C. elegans* embryo time-lapse with cell shapes optimized through FIDES

**Video 4:** Ablation of anterior zygote cortex during 70% polarization

**Video 5:** Ablation of posterior zygote cortex during 70% polarization

**Video 6:** Ablation of AB cortex

**Video 7:** Ablation of ABpr cortex

**Video 8:** Ablation of ABpl cortex

