## Supplementary Information for "Image-based force inference by biomechanical simulation"

#### 1 Supplementary text

##### 1.1 DCM force balance

Here, we provide the details of the deformable cell model, based on Cuvelier et al. [1]. Cells behave as pressurized liquid "bubbles" with surface tension, adhesive tension, and internal pressure. The actomyosin cortex of a cell is represented by a viscous shell which is approximated by a triangulated surface mesh with effective thickness $t_c$, and nodal positions $\mathbf{x}_i$. In the following sections we will discuss different force contributions and their implementation.

###### Surface tension

Tension generated in the cortex results from actomyosin contractility, which is represented as an effective surface tension $\gamma$ in the shell model. Adhesive tension $\omega_{ij}$ for contacting cells $i, j$ explains the reduction of contractility in interfacial surfaces, as well as the interfacial adhesion energy. Hence, the net surface tension $\tau$ of cell $i$ is given by

$$\tau_i = \begin{cases} \gamma_i & \text{for free surfaces} \\ \gamma_i - \omega_i & \text{for cell-cell contacting surfaces.} \end{cases}$$

The interfacial tension $\gamma_{ij}$ is the sum of the two contacting individual cortex tensions as

$$\gamma_{ij} = \gamma_i + \gamma_j - \omega_i - \omega_j = \gamma_i + \gamma_j - \omega_{ij}$$

where adhesive tension per cell is chosen such that the force at the cell-cell junction is also balanced at the level of the individual cell in the interface plane

$$\omega_i = \gamma_i \left(\cos(\theta_{iI}) + 1\right)$$
$$\omega_j = \gamma_j \left(\cos(\theta_{jI}) + 1\right).$$

We compute $\cos(\theta_{iI})$ and $\cos(\theta_{jI})$ from the balance of forces at the cell-cell junction

$$\gamma_i \cos(\theta_{iI}) + \gamma_j \cos(\theta_{jI}) + \gamma_{ij} = 0$$
$$\gamma_i \sin(\theta_{iI}) - \gamma_j \sin(\theta_{jI}) = 0.$$

We get

$$\cos(\theta_{iI}) = \frac{-\gamma_i^2 + \gamma_j^2 - \gamma_{ij}^2}{2\gamma_i\gamma_{ij}}$$
$$\cos(\theta_{jI}) = \frac{\gamma_i^2 - \gamma_j^2 - \gamma_{ij}^2}{2\gamma_j\gamma_{ij}}$$
$$\cos(\theta_{ij}) = \frac{-\gamma_i^2 - \gamma_j^2 + \gamma_{ij}^2}{2\gamma_i\gamma_j}.$$

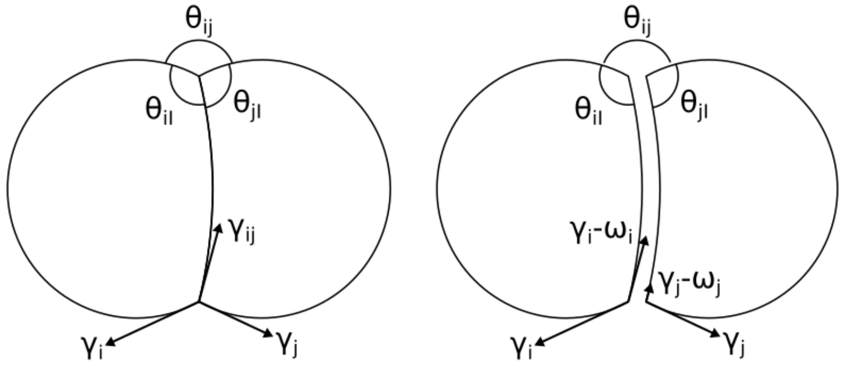


Figure S1: **Interfacial and adhesive tension at the cell-cell interface.** Where two cells connect, we consider adhesive tension $\omega_{ij}$ equivalent to interfacial tension $\gamma_{ij}$ when surface tensions $\gamma_i, \gamma_j$ at free surfaces are known.

Therefore, the free parameters in the system become $\gamma_i$, $\gamma_j$, and $\gamma_{ij}$ (or total adhesive tension $\omega_{ij} = \omega_i + \omega_j$ that we report in the manuscript). We have previously shown that our model predicts correct curvature and junction contact angles for varying adhesive tensions and surface tensions in Cuvelier et al. [1].

Using the derivation of Fedosov et al. [2], the direct force contribution of surface tension $\tau$ of node $i$ is given by

$$\mathbf{F}_i^{\text{act}} = \frac{\tau}{2}(\mathbf{x}_k - \mathbf{x}_j) \times \hat{\mathbf{n}}_A, \tag{1}$$

where $i, j, k$ are nodes of a single triangle $A = (ijk)$.

##### Internal pressure

Since the cytoplasm is assumed to be incompressible, active volume control is implemented via a cytoplasmic pressure. We assume the equilibrium volume $V_\alpha^*$ of a cell $\alpha$ is constant and implemented a proportional-integral (PI) pressure controller to enforce this. The cytoplasmic pressure $P_\alpha^{\text{cyt}}(t)$ due to the volume controller is then estimated as

$$P_\alpha^{\text{cyt}}(t) = -K\epsilon_\alpha(t) - \frac{K}{T_I}\int_0^t \mathrm{d}s\ \epsilon_\alpha(s), \tag{2}$$

for a cell with volumetric strain $\epsilon_\alpha(t) = (V_\alpha(t) - V_\alpha^*)/V_\alpha^*$. $T_I$ is typically chosen as one order of magnitude larger than the time-resolution with which the system is updated. The total pressure acting on a node

$$P_i(t) = \frac{2\gamma}{R_\alpha^{\text{cell}}} + P_\alpha^{\text{cyt}}(t),$$

includes an additional offset pressure $2\gamma/R_\alpha^{\text{cell}}$ to balance the actomyosin contractility in the cortex. This ensures cells are in mechanical equilibrium at the start of the simulations. The force acting on node $i$ due to nodal pressure $P_i(t)$, is obtained by integrating over the node associated oriented Voronoi area $S_i$. Given the pressure is constant over the cell surface, the force is simply given by

$$\mathbf{F}_i^{\text{cyt}}(t) = \mathbf{S}_i\, P_i(t). \tag{3}$$

##### Medium damping

A general drag force

$$\mathbf{F}_i^{\text{drag}} = -\Gamma_{ii}^{\text{f}} \cdot \mathbf{v}_i,$$

where

$$\Gamma_{ii}^{\text{f}} = \frac{3\eta^{\text{f}}}{2R^{\text{cell}}} S_i \mathbb{I}, \tag{4}$$

is included to account for the liquid drag between the cells and their medium due to the fluid viscosity $\eta^{\text{f}}$. When dealing with arbitrary shapes this approximation is no longer correct. Even though $F_i^{\text{drag}}$ is small compared to other dissipative forces, it is still used to improve the stability of the numerical integration scheme as it ensures the resistance matrix is positive definite, see further in equation of motion. [3]

##### Cortex viscosity

A much larger contribution to energy dissipation arises from cortex viscosity $\eta$. The viscous damping force between two connected nodes is given by

$$\mathbf{F}_{ij}^{\text{visc}} = \Gamma_{ij}^{c}(\mathbf{v}_j - \mathbf{v}_i), \tag{5}$$

with friction elements

$$\Gamma_{ij}^{\text{c}} = \frac{t_c \eta}{\sqrt{3}} \mathbf{I},$$

and $\mathbf{I}$ identity.

##### Wet contact friction

A viscous contact force is included to account for drag between contacting surfaces with friction constant $\xi$ (units of Pa·s/m), expressing the scaling between a dissipative traction $\mathbf{T}_c$ and the sliding velocity of two contacting surfaces $A$ and $B$

$$\mathbf{T}_{AB}^{\text{fric}} = -\xi \Delta \mathbf{v}_{AB}. \tag{6}$$

In the particle-based model, the contact drag force acting on node $i$ of triangle $A$ of the contact pair $(AB)$ is computed as

$$\mathbf{F}_{AB,i}^{\text{fric}} = \Gamma_{AB}^{\text{fric}} \cdot \sum_{k \in B} w_{ik}^{AB}\ (\mathbf{v}_k - \mathbf{v}_i) \tag{7}$$

determined by a friction tensor $\Gamma_{AB}^{\text{fric}}$ and weights $w_{ik}^{AB}$ per node $k$ of the $B$ triangle. $w_{ik}^{AB}$ are assumed to scale with the relative contribution of the nodal contact forces to the overall contact force thus

$$w_{ik}^{AB} = \frac{\left(\mathbf{F}_{AB,i}^{\text{adh}} + \mathbf{F}_{AB,k}^{\text{adh}}\right) \cdot \hat{\mathbf{n}}_{AB}}{6 \sum\limits_{\forall k \in B} \mathbf{F}_{AB,k}^{\text{adh}} \cdot \hat{\mathbf{n}}_{AB}}$$

is used as an approximation. $\Gamma^{\text{fric}}_{AB}$ for a given contact area $S_{AB}$ is estimated as

$$\Gamma^{\text{fric}}_{AB} = S_{AB}\left[\xi^{\perp}\hat{\mathbf{n}}_{AB}\otimes\hat{\mathbf{n}}_{AB} + \xi^{\parallel}\left(\mathbb{I} - \hat{\mathbf{n}}_{AB}\otimes\hat{\mathbf{n}}_{AB}\right)\right],$$

with normal and tangential friction coefficients $\xi^{\perp}$ and $\xi^{\parallel}$ respectively. Note that (7) can be equivalently formulated as

$$\mathbf{F}^{\text{fric}}_{AB,i} = \sum_{k\in B}\Gamma^{\text{fric}}_{AB,ik}\cdot\left(\mathbf{v}_k - \mathbf{v}_i\right),$$

if we denote

$$\Gamma^{\text{fric}}_{AB,ik} = w^{AB}_{ik}\Gamma^{\text{fric}}_{AB}. \tag{8}$$

#### Contact forces

In the mechanical cell model, we assume a fixed and uniform adhesive tension $\omega$ across the cell surface. The adhesive tension incorporates cell-cell adhesive interactions as well as the effect of cortical tension reduction at the cell-cell interface [4, 5, 6]. The total adhesive tension $\omega$ acting on interface $(AB)$ is given by $\omega = \omega_A + \omega_B$. A linear force, scaling with contact overlap $\delta$, was introduced to represent repulsive interactions between surfaces and ensure stable contacts. The contact potential for contact-pair $(AB)$ with contact area $S_{AB}$ is then given by

$$E^{\text{adh}}_{AB} = \left(\frac{\delta_{AB}}{h_0} - \frac{\delta^2_{AB}}{2h^2_0}\right)\omega S_{AB}$$

where a distinction can be made between the attractive and repulsive energy contributions respectively. The effective range of adhesion $h_0$ was incorporated by virtually translating contact surfaces along their normal direction. The contact overlap distance $\delta_{AB}$ is calculated with respect to these translated surfaces. The resulting contact pressure $P^{\text{adh}}_{AB}$ is estimated with a linear contact pressure model where no energy is stored in deformation at equilibrium

$$P^{\text{adh}}_{AB}\left(\mathbf{x}\right) = k_{AB}\delta_{AB}\left(\mathbf{x}\right) - P^0_{AB},$$

contact stiffness $k_{AB}$ and adhesive pressure $P^0_{AB}$ need to be set. To ensure that at equilibrium ($P^{\text{adh}}_{AB}\equiv 0$) the work of adhesion $\omega$ is recovered, contact stiffness has to be defined as $k_{AB} = P^0_{AB}/h_0$ with $P^0_{AB} = \omega/h_0$. The specific value $h_0$ does not influence simulation outcome due to the scale separation, $h_0 \ll R^{\text{cell}}$. This is valid for all performed simulations as the effective range of adhesion $h_0$ was estimated as 300 nm. [7] Contact overlap distance at point $\mathbf{x}$ was defined as

$$\delta_{AB}\left(\mathbf{x}\right) = \max\left\{0;\ 2\left(\mathbf{x} - \mathbf{s}_{AB}\right)\cdot\left(\hat{\mathbf{n}}_{AB}\times\hat{\mathbf{l}}_{AB}\right)\tan\theta\right\},$$

where $\theta$ is the contact angle, $\hat{\mathbf{n}}_{AB}\propto\left(\hat{\mathbf{n}}_B - \hat{\mathbf{n}}_A\right)\times\hat{\mathbf{l}}_{AB}$ the contact normal and $\mathbf{s}_{AB}$ is an arbitrary point on the intersection line defined by $\hat{\mathbf{l}}_{AB}\propto\hat{\mathbf{n}}_A\times\hat{\mathbf{n}}_B$, as reported by Smeets et al. [8] The resulting contact forces and moment are obtained by integrating the total contact pressure over the oriented contact area $\mathbf{S}_{AB}$,

$$\mathbf{F}^{\text{adh}}_{AB} = \int \mathrm{d}\mathbf{S}_{AB}(\mathbf{x})\ P^{\text{adh}}_{AB}(\mathbf{x}). \tag{9}$$

To distribute the contact forces acting on the contact plains to the nodal contact forces $\mathbf{F}^{\text{adh}}_{AB,i}$, it is assumed that the nodal forces are collinear with the contact normal $\hat{\mathbf{n}}_{AB}$. This results in a system of linear equations per contact pair $(AB)$

$$\sum_{i\in A} \mathbf{F}^{\mathrm{adh}}_{AB,i} = -\int\limits_{\mathbf{x}\in A\cap B} \mathrm{d}\mathbf{S}_{AB}(\mathbf{x})\; P^{\mathrm{adh}}_{AB}(\mathbf{x})$$

$$\sum_{i\in A} \left[(\mathbb{I} - \hat{\mathbf{n}}_{AB}\otimes\hat{\mathbf{n}}_{AB})\cdot(\mathbf{x}_i - \mathbf{x}_{AB})\right]\times\mathbf{F}^{\mathrm{adh}}_{AB,i} = \int\limits_{\mathbf{x}\in A\cap B} \mathrm{d}\mathbf{S}_{AB}(\mathbf{x})\times(\mathbf{x}-\mathbf{x}_{AB})\; P^{\mathrm{adh}}_{AB}(\mathbf{x}) \tag{10}$$

which guarantees a unique solution for every $\mathbf{F}^{\mathrm{adh}}_{AB,i}$. The contact point $\mathbf{x}_{AB}$ is defined as the geometric center of the intersection polygon $A\cap B$. [8] The integrals on the right-hand side are calculated by using a numerical quadrature rule presented in [9] with 7 quadrature points covering each triangle of the triangulated intersection polygon $A\cap B$ obtained as an intersection of projected triangles on a common plane. The precise choice of the number of quadrature points has no impact on the results in the simulation, as long as adequate quadrature coordinates and weighting values are chosen. The solution for this system is presented and discussed in the work of Smeets et al. [8].

This same contact model is used for cell-eggshell contacts where we assume low adhesive tension ($\omega_h$), and high contact stiffness ($k_{AB} = \frac{1}{3}10^5\frac{2\gamma}{R_0^2}$). The eggshell itself is assumed undeformable, meaning we don't integrate the node positions.

#### Protrusion forces

The protrusion model is defined by two spherical coordinates directing $\mathbf{p}_p$, an angle $\alpha$ defining the protrusion width and a total protruding force $F_p$. We model a protrusive pressure in direction $\mathbf{p}_p$ per node $i$ at the front end of the cell by the difference of Gaussian kernels as

$$P_i^P = \beta_P\left(2\exp\frac{-\theta_i^2}{2\alpha^2} - \exp\frac{-\theta_i^2}{8\alpha^2}\right). \tag{11}$$

Here $\beta_P$ is the scaling factor determining the total protrusive force $F_P$, $\alpha$ gives a 3D angle indicating the width of protrusion pressure (fig. S2), and $\theta_i$ is the 3D angle for every node between the line from the cell center $x_c$ in direction $\mathbf{p}_p$ and the line $\mathbf{x}_i - \mathbf{x_c}$ such that

$$\cos\theta_i = \frac{\mathbf{p}_p\cdot(\mathbf{x}_i-\mathbf{x}_c)}{||\mathbf{p}_p||\;||(\mathbf{x}_i-\mathbf{x}_c)||}. \tag{12}$$

We set $\beta_P$ such that the total pushing force in direction $\mathbf{p}_p$ becomes

$$F_p = \int P_i^P H(P_i^P)\mathbf{p}_p\cdot d\mathbf{S}_i, \tag{13}$$

with the Heaviside function

$$H(P_i^P) = \begin{cases} 1\,, & P_i^P >= 0 \\ 0\,, & P_i^P < 0 \end{cases} \tag{14}$$

To make sure that the net force applied from protrusion is zero, we add a reaction pressure

$$P_i^R = \beta_R\exp\frac{-\theta_i^2}{8\alpha^2} \tag{15}$$

and set the reaction pressure scale $\beta_R$ such that

$$0 = \int \left(P_i^P + P_i^R\right) \mathbf{p}_p \cdot d\mathbf{S}_i. \tag{16}$$

The force from cell protrusion per node $i$ is computed as

$$\mathbf{F}_i^P = \mathbf{S}_i P_i^P. \tag{17}$$

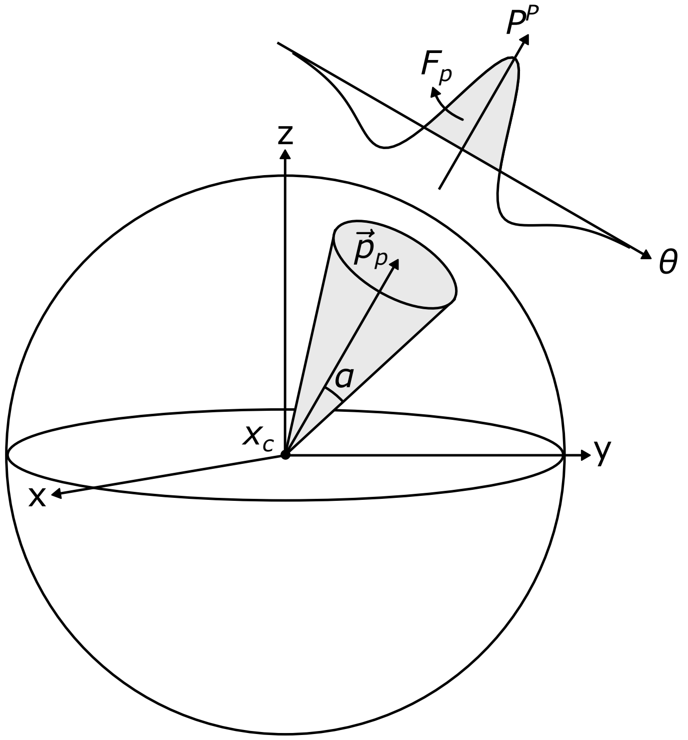


Figure S2: **Protrusion model.** The protrusion is characterized by an orientation vector $\mathbf{p}_p$, angle $\alpha$, and total protruding force $F_p$.

##### Cytokinetic ring forces

The cytokinetic ring model is defined by two spherical coordinates directing $\mathbf{p}_R$, spring constant $k$, and an offset distance $d$. The cell division plane is defined through point $\mathbf{x}_c + d\,\mathbf{p}_R$ with orientation $\mathbf{p}_R$, giving it an offset from cell center $x_c$. We add spring forces between the connecting nodes $i, j$ of the contractile ring as

$$\mathbf{F}_i^R = k\left(1 - \frac{\Delta x_0}{\Delta x_R}\right)(\mathbf{x}_j - \mathbf{x}_i) \tag{18}$$

where $k$ is the effective spring stiffness, $\Delta x_0$ is the constant reference length of the ring chosen as 20µm, and $\Delta x_R$ is the instantaneous length of the contractile ring. Furthermore, since we observe in vivo that cytokinesis gives rise to a smooth contractile band, we add a smoothing force acting on ring nodes $i$ and neighboring nodes $j$ as

$$\mathbf{F}_j^{R,S} = \mathbf{p}_i \frac{k}{c}\left[\mathbf{p}_i \cdot (\mathbf{x}_i - \mathbf{x}_j)\right] \tag{19}$$

with c set to 5m. To make sure that we have a net-zero force balance

$$\mathbf{F}_i^{R,S} = -\mathbf{F}_j^{R,S} \tag{20}$$

with $\mathbf{p}_i = (\mathbf{x}_i - \mathbf{x}_c)/||\mathbf{x}_i - \mathbf{x}_c||$ the direction from the cell center to the ring node.

To keep the volumes of emerging daughter cells stable, we add pressure per daughter based on surface area deviation

$$P^D = K_D(S_D^* - S_D(t)) \tag{21}$$

with $K_D$ a proportional controller constant ($1 \times 10^{12}\,\mathrm{Pa\,m^{-2}}$), $S_D^*$ the equilibrium area and $S_D(t)$ the instantaneous area. The force per daughter node $i$ becomes

$$\mathbf{F}_i^D = \mathbf{S}_i P_i^D. \tag{22}$$

##### Equation of motion

In the overdamped cellular environment, inertial forces may be neglected [10]. Based on the different contributions described above, the force balance for node $i$ gives

$$\begin{aligned}\mathbf{F}_i^{\mathrm{act}} + \mathbf{F}_i^{\mathrm{cyt}} + \mathbf{F}_i^{P} + \mathbf{F}_i^{R} + \mathbf{F}_i^{R,S} + \mathbf{F}_i^{D} + \sum_{(AB):i\in A} \mathbf{F}_{AB,i}^{\mathrm{adh}} \\ = \Gamma_{ii}^{\mathrm{f}} \cdot \mathbf{v}_i + \sum_j \Gamma_{ij}^{\mathrm{c}} \cdot (\mathbf{v}_i - \mathbf{v}_j) + \sum_{(AB):i\in A} \sum_{k\in B} \Gamma_{AB,ik}^{\mathrm{fric}} \cdot (\mathbf{v}_i - \mathbf{v}_k)\end{aligned}$$

which can be abbreviated as

$$\mathbf{F}_i = \sum_j \Gamma_{ij} \cdot \mathbf{v}_j \tag{23}$$

for a system consisting of $N$ nodes, where

$$\Gamma_{ij} = \begin{cases} \Gamma_{ii}^{\mathrm{f}} + \sum_{k:k\neq i} \Gamma_{ik}^{\mathrm{c}} + \sum_{(AB):i\in A} \sum_{k\in B} \Gamma_{AB,ik}^{\mathrm{fric}}, & i = j, \\ -\Gamma_{ij}^{\mathrm{c}} - \sum_{(AB):i\in A, j\in B} \Gamma_{AB,ij}^{\mathrm{fric}}, & i \neq j. \end{cases}$$

The Cartesian components of the overall force vectors can be represented as a single $(3N \times 1)$ column matrix, while the friction matrices can be assembled to a single $(3N \times 3N)$ sparse symmetric positive definite friction matrix [11]. The conjugate gradient method (CGM) is used to efficiently solve the system for the node velocities $\{\mathbf{v}_j\}$ at each iteration. Positions of the nodes are subsequently updated using a semi-implicit Euler integration scheme.

$$\mathbf{x}_i(t + \Delta t) = \mathbf{x}_i(t) + \Delta t\, \mathbf{v}_i(t + \Delta t), \tag{24}$$

where $\mathbf{v}_i(t + \Delta t)$ are the projected new velocities obtained by solving eq. 23 at time $t$ via the CGM.

#### 1.2 Protein signal correction

First, the raw protein signal from a microscopy time-lapse is smoothed using a Gaussian kernel with a sigma of 0.2 microns. The smoothed signal is then projected onto the cell meshes, where each mesh node takes on the value of the nearest pixel.

To account for bleaching over time and signal weakening in z, a linear generalized additive model (GAM) is applied. Time (t) and z are fitted to piecewise second-order splines using the following pyGAM syntax:

```
gam = LinearGAM(s('t', n_splines=10, spline_order=2)
              + s('z', n_splines=20, spline_order=2))
```

The signal is subsequently normalized to a range of 0 to 1.

For calculating the protein value at a cell-cell interface or for an entire cell, a simple average of the corresponding mesh nodes is used.

##### 1.3 Automatic detection of protrusions and cytokinetic rings

Both rings and protrusions can be detected and assigned automatically, based on a quantitative analysis of the cell shape. To demonstrate this, we used FlowShape [12] to set up a pipeline for the detection of protrusions and cytokinetic rings, along with their orientation. We selected a representative set of 31 cell shapes, which were then manually labeled with the number of protrusions. Further, a cell was labeled as having a cytokinetic ring when it has divided in the next frame in the time-lapse.

The protrusion detector is a Laplacian of Gaussian filter to detect regions of high curvature, followed by a simple threshold (the same filter as described in van Bavel et al. [12]). This design reached an accuracy of 90% on our dataset. To detect cytokinetic rings, we made a synthetic 'template' of a dividing cell, and try to match it to all cells by maximizing the cross-correlation. Using a single template of a symmetric division, we managed to match all but one cell, giving an accuracy of 97%. The missed cell shows a highly asymmetric division, and could additionally be correctly classified by introducing a second template that accounts for asymmetry. The matched templates also contain information about the orientation of the division plane, but we did not quantify these. Code and data for this pipeline can be found in the FlowShape repository: `https://bitbucket.org/pgmsembryogenesis/flowshape/src/main/demo/detect_features.ipynb?viewer=nbviewer`.

#### 2 Supplementary figures

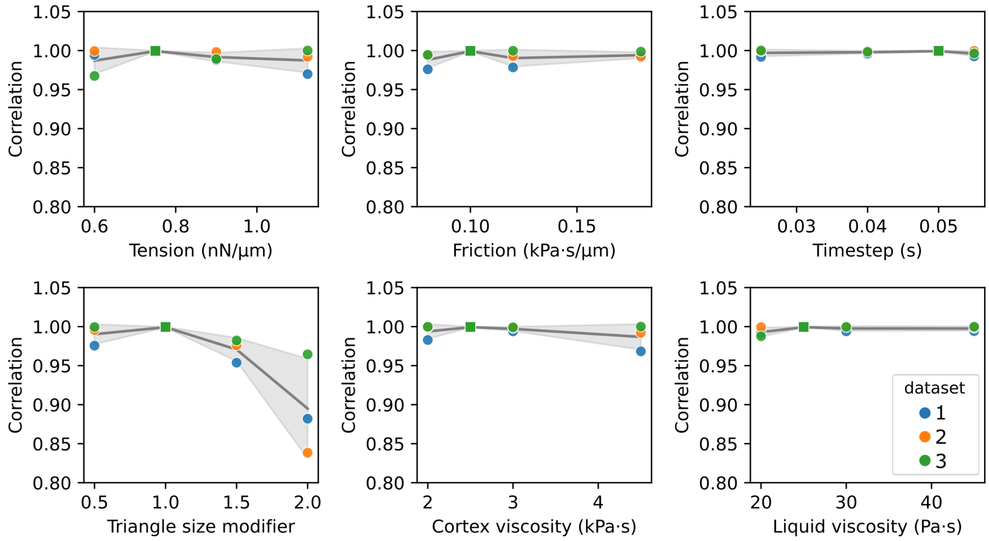


Figure S3: **Sensitivity analysis for FIDES physical parameters.** To test the effect of the initial parameters on the FIDES inferences, we performed a sensitivity analysis for three different embryos. Scores are the correlation coefficient of surface and adhesive tension results, compared to a baseline result. First, the original experiment was repeated threefold and the results were averaged to make this baseline. The repeats were very consistent, each scoring near perfect correlation to the baseline result. We then repeated the experiment, having varied the initial surface tension, contact friction, simulation timestep, cortex viscosity, or liquid viscosity as simple static parameters to FIDES. We also tried different mesh sizes, modifying the triangle size during PiCS preprocessing to yield a different input mesh for FIDES. This reveals that our inferences are quite robust against a slight change in physical parameters. However, when the triangle size is increased too much, a lot of contact detail gets lost and the inferences start to diverge. This follows our expectations, as the triangle size was chosen to keep enough detail, without being unnecessarily high, which would hurt simulation time. The gray band represents the data range between the three tested embryos.

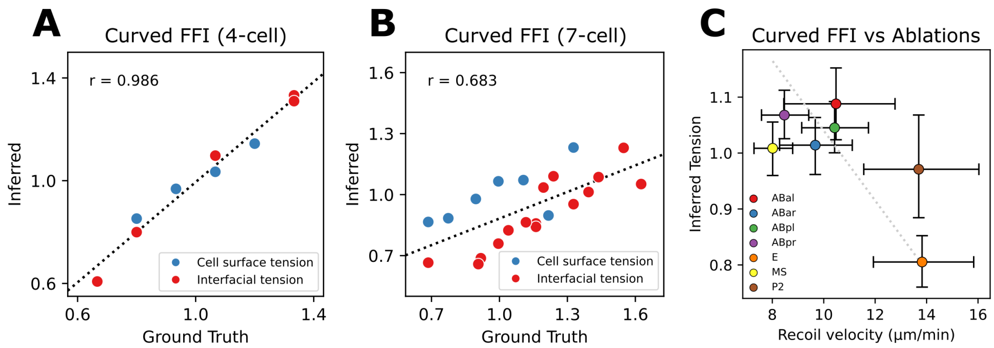


Figure S4: **Synthetic and ablation performance for curved FFI.** A: Performance for the simple 4-cell embryo. B: Performance for the advanced 7-cell embryo. C: Comparison of curved FFI surface tensions with ablation velocities.

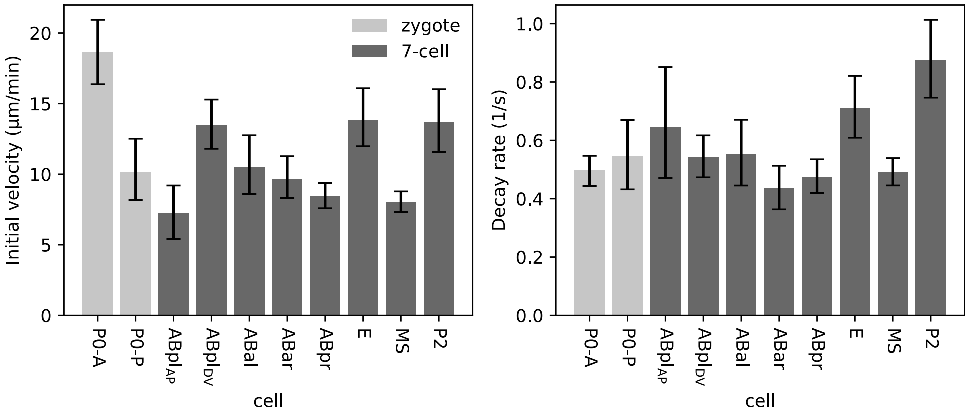


Figure S5: **Laser ablation exponential results.** Exponential parameters for the zygote (P0) and 7-cell cuts. 95% confidence intervals are calculated using bootstrap (1000 samples).

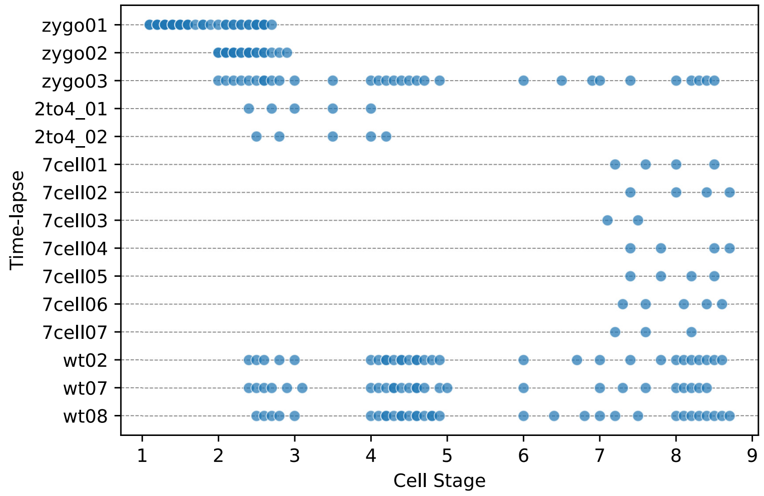


Figure S6: **Embryo time-lapses and their coverage of early development.**

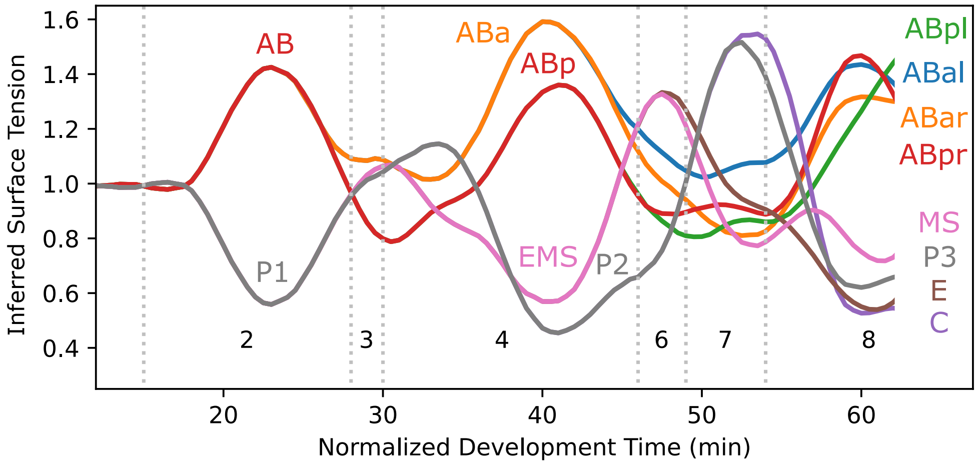


Figure S7: **Timeline of inferred FIDES surface tensions between 1- and 8-cell stage.** The inferred surface tensions for all embryos are aggregated for each cell and smoothed over time. Surface tensions are always normalized to average 1, equalizing their levels over time. Relative differences can be observed at any timepoint, but the evolution of real average tension is unknown.

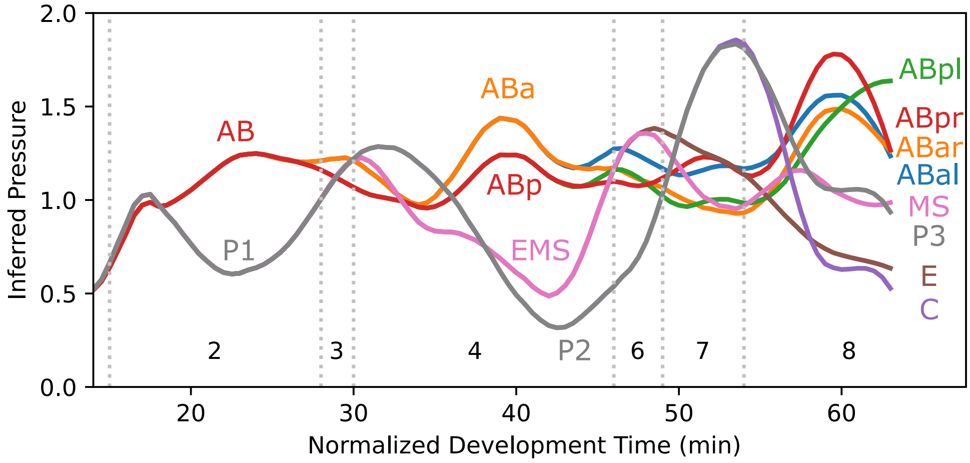


Figure S8: **Timeline of inferred FIDES cell pressures between 1- and 8-cell stage.** The inferred pressures for all embryos are aggregated for each cell and smoothed over time. These values are mostly dependent on the surface tension for the corresponding cell, visible in fig. S7. Relative differences can be observed at any timepoint, but the evolution of real pressure is unknown. For that reason, we further normalized the pressures by dividing with reference pressure $2\gamma_0/R_0$.

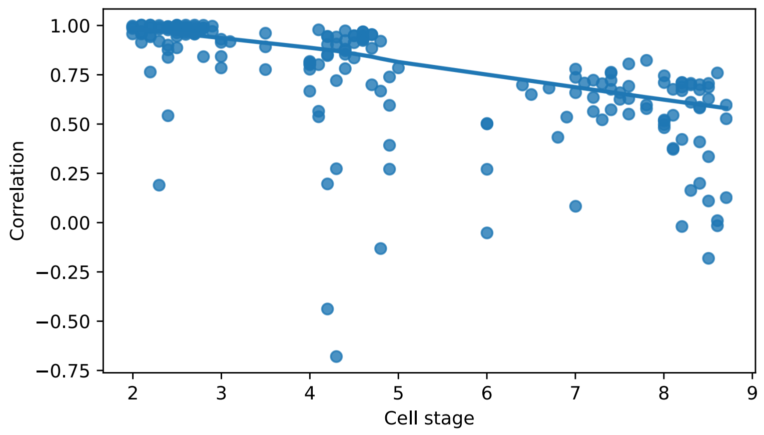


Figure S9: **Correlation between results from FIDES and tangent FFI.** For every embryo, the Pearson's correlation coefficient between the two methods is plotted, considering both inferred surface and adhesive tension values. The lowess fit was used to observe a decreasing trend. Both methods are in strong agreement for the early cell stages, as the cells remain similar to a foam. From the 6-cell stage onwards, the inferences from both methods diverge as the assumptions of foam models are less satisfied.

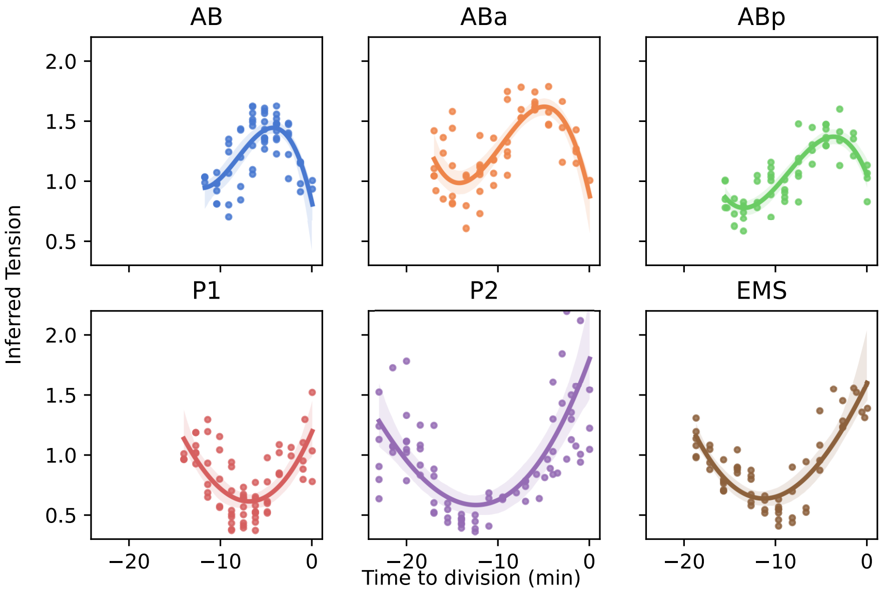


Figure S10: **Surface tension inferences during mitotic rounding.** Scatterplot of the individual tension inferences leading up to cell division. Surface tensions within a sample are always normalized to average 1, so these plots only describe how one surface tension evolves compared to the average. We fit a third order polynomial regression with the band representing a 95% confidence interval of the regression estimate.

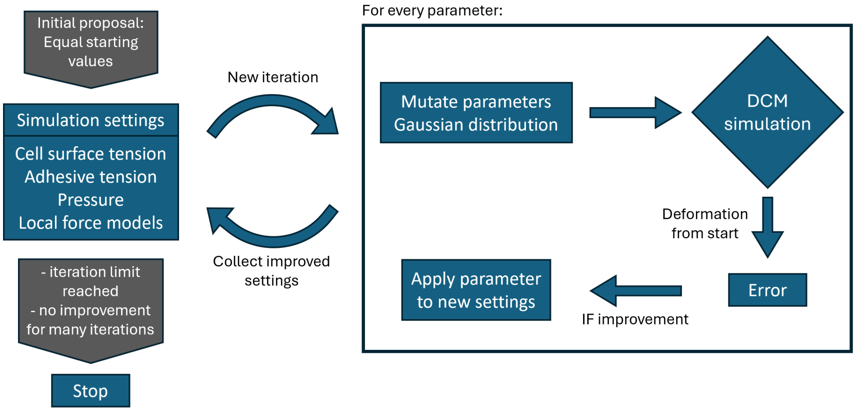


Figure S11: **Parameter optimization strategy for FIDES.** Repeated iterations evaluate parameter changes by computing the deformation error in a DCM simulation. Gaussian learning rate and bounds for all parameters can be found in the code (`solve_shape.py`).

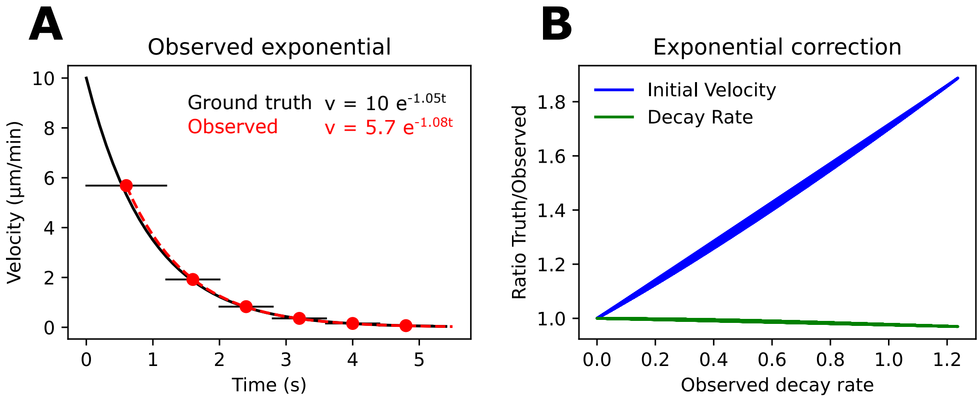


Figure S12: **Correction for laser ablation velocity.** A: Simulation of ablation observation shows that the initial velocity is underestimated, especially when the exponential decay is fast. B: Performing this simulation 1000 times with different exponential parameters, the initial recoil velocity needs a correction that scales linearly with the observed decay rate, while the decay rate is minimally affected.

#### 3 Supplementary tables

| Condition | Velocity | Decay | Experiments | Markers |
|---|---|---|---|---|
| P0-A | 18.7 | 0.50 | 19 | 1498 |
| P0-P | 10.1 | 0.54 | 14 | 738 |
| ABal | 10.5 | 0.55 | 7 | 420 |
| ABar | 9.7 | 0.43 | 11 | 602 |
| $\mathrm{ABpl_{DV}}$ | 13.5 | 0.47 | 18 | 1137 |
| $\mathrm{ABpl_{AP}}$ | 7.2 | 0.64 | 7 | 296 |
| ABpr | 8.5 | 0.48 | 21 | 1098 |
| E | 13.9 | 0.71 | 21 | 1175 |
| MS | 8.0 | 0.49 | 21 | 1332 |
| P2 | 13.7 | 0.87 | 14 | 794 |

Table S1: **Numeric ablation results and sample sizes.**

| Element | V | P | R | Deviation (°) | Total force (µN) | N |
|---|---|---|---|---|---|---|
| P0 ring | 0.00 | 1.00 | −0.02 | 0 | $0.008 \pm 0.007$ | 3 |
| AB ring | −0.86 | 0.51 | 0.00 | 12 | $0.76 \pm 0.14$ | 7 |
| P1 ring | −0.02 | 0.99 | −0.12 | 4 | $0.29 \pm 0.17$ | 8 |
| ABa ring | −0.53 | 0.07 | 0.85 | 5 | $0.46 \pm 0.02$ | 5 |
| ABp ring | −0.38 | 0.21 | 0.9 | 10 | $0.27 \pm 0.16$ | 7 |
| EMS ring | 0.10 | 0.95 | −0.28 | 3 | $0.26 \pm 0.03$ | 3 |
| P2 ring | 0.93 | 0.37 | −0.01 | 11 | $0.36 \pm 0.17$ | 9 |
| ABpl ventral protrusion | 0.96 | 0.13 | 0.23 | 10 | $0.0034 \pm 0.0025$ | 24 |
| ABpl anterior protrusion | −0.23 | −0.97 | 0.07 | 10 | $0.0032 \pm 0.0017$ | 24 |
| ABpl dorsal lamellipodium | −0.93 | −0.36 | 0.09 | 14 | $0.0018 \pm 0.0019$ | 24 |
| ABpr ventral protrusion | 0.99 | −0.10 | −0.13 | 9 | $0.0032 \pm 0.0021$ | 24 |

Table S2: **Averaged parameters of local force generating activities.** The 3D coordinate system is aligned to the embryo symmetry axes, with V representing the ventral side of the DV axis, P the posterior side of the AP axis, and R the right side of the LR axis. For cytokinetic rings, only data from the last 2 minutes before division is used. The orientation of a ring is a unit vector perpendicular to the division plane, while the total force on the ring is defined as spring constant $k$, multiplied by ring length. Local protrusions are averaged over the entire 7-cell stage. The total force is the pushing force on the protrusion, while the orientation is given as a unit vector pointing from the cell center to the protrusion tip. For all features, the orientation is a weighted average of the individual orientations, using the total force they exerted as weights. The variation of orientations is given as an angular weighted standard deviation, using the same weights.

| Parameters | Symbol | Synthetic | PiCS | FIDES | Units | Sources |
|---|---|---|---|---|---|---|
| Surface tension | $\gamma$ | 0.75 | 1.5 | 0.75 | $\mathrm{nN\,\mu m^{-1}}$ | [13, 1] |
| Adhesion cell-cell | $\omega_{ij}$ | 0.75 | 0.5 | 0.5 | $\mathrm{nN\,\mu m^{-1}}$ | [11, 13] |
| Adhesion cell-hull | $\omega_h$ | 0.01 | 0.01 | 0.01 | $\mathrm{nN\,\mu m^{-1}}$ | assumed minimal |
| Bulk modulus cell | $K$ | 1.5 | 1 | 2 | kPa | [1] |
| K integrative term | $T_I$ | $2\Delta t$ | $2\Delta t$ | $2\Delta t$ | s | |
| Thickness cortex | $t_c$ | 300 | 300 | 300 | µm | [14, 1] |
| Adhesive range | $h$ | 200 | 250 | 200 | nm | |
| Normal Friction | $\xi^{\perp}$ | 1 | 1 | 1 | $\mathrm{kPa\,s\,\mu m^{-1}}$ | |
| Tangential Friction | $\xi^{\parallel}$ | 0.05 | 0.01 | 0.1 | $\mathrm{kPa\,s\,\mu m^{-1}}$ | |
| Liquid viscosity | $\eta^{\mathrm{f}}$ | 25 | 500 | 25 | Pa s | [13] |
| Cortex viscosity | $\eta$ | 2 | 2 | 2.5 | kPa s | |
| Cell radius | $R_0$ | 10 | 10 | 10 | µm | |
| Simulation timestep | $\Delta t$ | 0.05 | 0.04 | 0.05 | s | stable |
| Simulation duration | | 150 | 1.5 | 2.5 | s | |

Table S3: **Mpacts physical parameters.** The synthetic parameters are used to generate synthetic embryos. PiCS parameters are used for the final relaxation step of PiCS image segmentation. FIDES parameters are used for the stability tests of input meshes.

### 4 Supplementary videos

**Video 1–3:** *C. elegans* embryo time-lapse with cell shapes optimized through FIDES (dataset wt02, wt07, and wt08 respectively). Arrows indicate local force generation. Colors are used to mark the different cells.

**Video 4:** Ablation of anterior zygote cortex during 70% polarization (P0-A, sample 220627-E3). F-actin is shown in red, NMY-2 in green, the image is 26 µm wide and frame time is 1.03 s.

**Video 5:** Ablation of posterior zygote cortex during 70% polarization (P0-P, sample 220914-E2). F-actin is shown in red, NMY-2 in green, the image is 26 µm wide and frame time is 1.05 s.

**Video 6:** Ablation of AB cortex (sample 220726-E6). F-actin is shown in red, NMY-2 in green, the image is 26 µm wide and frame time is 1.36 s.

**Video 7:** Ablation of ABpr cortex (sample 220919-E11). F-actin is shown in red, NMY-2 in green, the image is 26 µm wide and frame time is 1.36 s.

**Video 8:** Ablation of ABpl cortex (sample 220927-E3). F-actin is shown in red, NMY-2 in green, the image is 26 µm wide and frame time is 0.82 s.

1367-4811. doi: 10.1093/bioinformatics/btad383. URL http://dx.doi.org/10.1093/bioinformatics/btad383.

[13] Maxim Cuvelier, Jiří Pešek, Ioannis Papantoniou, Herman Ramon, and Bart Smeets. Distribution and propagation of mechanical stress in simulated structurally heterogeneous tissue spheroids. *Soft Matter*, 17(27):6603–6615, 2021. doi: 10.1039/d0sm02033h. URL https://doi.org/10.1039/d0sm02033h.

[14] Bart Smeets, Maxim Cuvelier, Jiri Pešek, and Herman Ramon. The effect of cortical elasticity and active tension on cell adhesion mechanics. *Biophysical Journal*, 116(5): 930–937, March 2019. doi: 10.1016/j.bpj.2019.01.015. URL https://doi.org/10.1016/j.bpj.2019.01.015.
